# Resolving temporal threat uncertainty by observational learning involves the amygdala, hippocampus and anterior insula

**DOI:** 10.1101/2025.10.21.682164

**Authors:** Simon Knobloch, Theresa Jansen, Lara Hille, Madeleine Müller, Leonie Rumpf, Christian Bauer, Jan Haaker

## Abstract

By observing others’ experiences, humans learn about threats while avoiding firsthand dangers. Yet, previous neuroscience research has focused on observational threats that are predictable. In firsthand learning, temporally predictable and unpredictable threats can be mechanistically discerned. In this study, we developed a novel observational paradigm in which participants learned from predictable and unpredictable observational threats, as well as a no-threat condition. Participants encountered the same conditions during an expression phase to investigate how the brain encodes predictive and unpredictive threat cues observed in others. Participants in Experiment 1 (n = 20, 14 female, 6 male) and Experiment 2 (n = 23, 9 female, 1 non-binary, 13 male) successfully learned threat contingencies, showing heightened threat expectations for predictive cues and unpredictable condition-onsets. Behavioral findings converged with neural (fMRI, Experiment 2) responses in the anterior insula during the expression phase. Reflecting the dynamic process of learning, amygdala activation in response to predictive threat cues displayed a linear decrease across trials. Interestingly, BOLD responses to others’ pain that was predictable, were enhanced within the amygdala, insula and hippocampus, compared to unpredictable conditions. Our findings suggest that humans can learn to resolve temporal threat uncertainty, through the observation of others. The present work thereby contributes to understanding the social aspects of fear and anxiety disorders.

## 1. Introduction

Accurate prediction of threats is key to survival. Since we live in sociocultural environments, observational threat learning enables us to predict dangers without direct encounters of threats (Berger, 1962; Olsson & Phelps, 2004; Vaughan & Lanzetta, 1980). In observational threat learning, the demonstrator’s expression of pain serves as an unconditioned aversive stimulus that enables learning of predictive threat cues (Olsson et al., 2007). Previous research has demonstrated similarities between observational and firsthand learning, reflected in comparable behavioral, psychophysiological, and neural responses. (Haaker, Yi, et al., 2017; Lindstrom et al., 2018; Olsson et al., 2020; Olsson et al., 2007).

In these protocols of observational threat learning, the observed US was predictable by non-social cues (e.g., geometric shapes predicting the observed US). Yet, from a functional perspective, humans have to learn about both, predictable threats and threats that occur unpredictably (Schmitz & Grillon, 2012). It remains, however, unclear, how unpredictable threats are learned by observation, i.e. in the absence of non-social predictors.

In firsthand learning, predictable and unpredictable threats result in phasic and sustained defensive responses, respectively (Schmitz & Grillon, 2012). When threats occur predictably (i.e. in association with cues), a phasic anxious arousal response is elicited in participants, as measured by skin-conductance responses or eye-blink startle, to the predictive cue. In contrast, when threats occur unpredictably (i.e. independent of cues), responses are characterized by sustained elevated arousal across the condition as a whole (Alvarez et al., 2011; Schmitz & Grillon, 2012). As such, predictable threats result in expectation of aversive outcomes when a signaling cue is present (cue conditioning), whereas unpredictable threats elevate expectation across the condition (context conditioning), relative to the cue. It has been argued that these two defensive reactions are not only discernable experimentally, but map well onto the distinct factors of different internalizing disorders (Schmitz & Grillon, 2012).

At the neural level, the amygdala and anterior insula play a central role in both firsthand (Buchel, Holmes, et al., 1998; Buchel, Morris, et al., 1998; Marschner et al., 2008; Quirk et al., 1995) and observational threat learning (Haaker, Yi, et al., 2017; Olsson et al., 2007). In firsthand learning, the hippocampus (Alvarez et al., 2008; Marschner et al., 2008) is central to learning about unpredictable threats, likely with involvement of the amygdala (Alvarez et al., 2008; Figel et al., 2019). Furthermore, the insular cortex has been shown to be involved in the anticipation of unpredictable threats (Alvarez et al., 2011; Geng et al., 2018; Herrmann et al., 2016; Hur et al., 2020; Radoman et al., 2021; Shankman et al., 2014). Taken together, although the learning of unpredictable threats through observation has not yet been thoroughly examined, the amygdala, insular cortex, and hippocampus are likely to play key roles in mediating learning about unpredictable aversive experiences.

In this study, we aimed to examine observational learning of predictable and unpredictable threats, where a demonstrator, viewed on screen by participants, encountered painful threat. This observed painful threat occurred either temporally predictably (predictable condition [P]) or temporally unpredictably (unpredictable condition [U]), compared to a no-pain condition [N]. We hypothesized (in Experiment 1) that participants are able to resolve the threat-related information solely through observation. We further hypothesized that participants would exhibit discriminable threat expectations, with higher threat expectancy for predictive cues and unpredictable conditions, compared with cues and conditions where no painful outcomes were observed. In Experiment 2, we aimed to replicate these behavioral findings using pre-registered, pre-defined, directional contrasts (e.g., higher cue prominence in predictable, compared to unpredictable conditions). In addition, we aimed to characterize the neural responses underlying learning predictable and unpredictable threats from others using the same pre-registered, pre-defined contrasts.

## 2. Methods

### 2.1 Paradigm

The conceptual basis for our observational learning adaptations was a well-known two-phase experimental paradigm, in which participants learn threat/cue contingencies by observation in stage one (observation phase), and in stage two (expression phase) directly encounter the cues learned to predict threat or safety (Haaker, Golkar, et al., 2017). Our current study adapted this protocol to examine observational learning of temporally predictable and unpredictable threats in comparison with a no threat condition. Specifically, only in P were participants able to learn when the observed pain would occur within individual trials, whereas in the U, observed pain occurred at temporally unpredictable time points. The paradigm therefore shares features with the established NPU paradigm for firsthand threat learning (Schmitz & Grillon, 2012). However, in contrast to the original NPU paradigm, our task was uninstructed, i.e. participants were not informed about the threat contingencies in each condition.

We report two experiments in this publication. In Experiment 1, we developed and tested an observational learning paradigm in a behavioral laboratory setting. In Experiment 2, we adapted the paradigm for use with fMRI. This enabled preregistration and replication of the behavioral findings from Experiment 1 in an independent sample in Experiment 2, while additionally examining the underlying neural correlates of observational threat learning. The conducted analyses were preregistered prior to data-analysis (Knobloch et al., 2024).

### 2.2 Experiment 1

#### 2.2.1 Participants and procedure

20 healthy participants (see Table 1 for descriptives) were recruited by online and black-board postings at the University Medical Center Hamburg. All participants were fluent in German and able to understand the task instructions. Further inclusion criteria were an age between 18 and 40, sufficient general health and normal or corrected-to-normal vision. Exclusion criteria were current or previous psychiatric disorder, current use of psychoactive medication or substances and current or previous neurological disease. Participants that had already participated in threat-conditioning studies at our institute within three months preceding the participation were also excluded. Eligibility criteria were assessed in structured interviews via telephone and verified on site by a physician. Study data was collected in a single appointment of approximately 60 minutes. After giving informed consent to participate in the study, participants filled out a questionnaire collecting sociodemographic data. The experiment consisted of two blocks of 25 minutes (see 2.2.2 Experimental design) and was conducted on a computer in the behavioral lab. After completing the experimental paradigm, participants filled in a post-experimental questionnaire, in which they answered questions about the cue/pain contingencies and rated the observed pain expression with regard to how natural they perceived the demonstrator’s pain expression to be and their empathy towards the demonstrator. All questionnaires were conducted via soSci (soscisurvey.de, München, Germany).

**Table 1.**
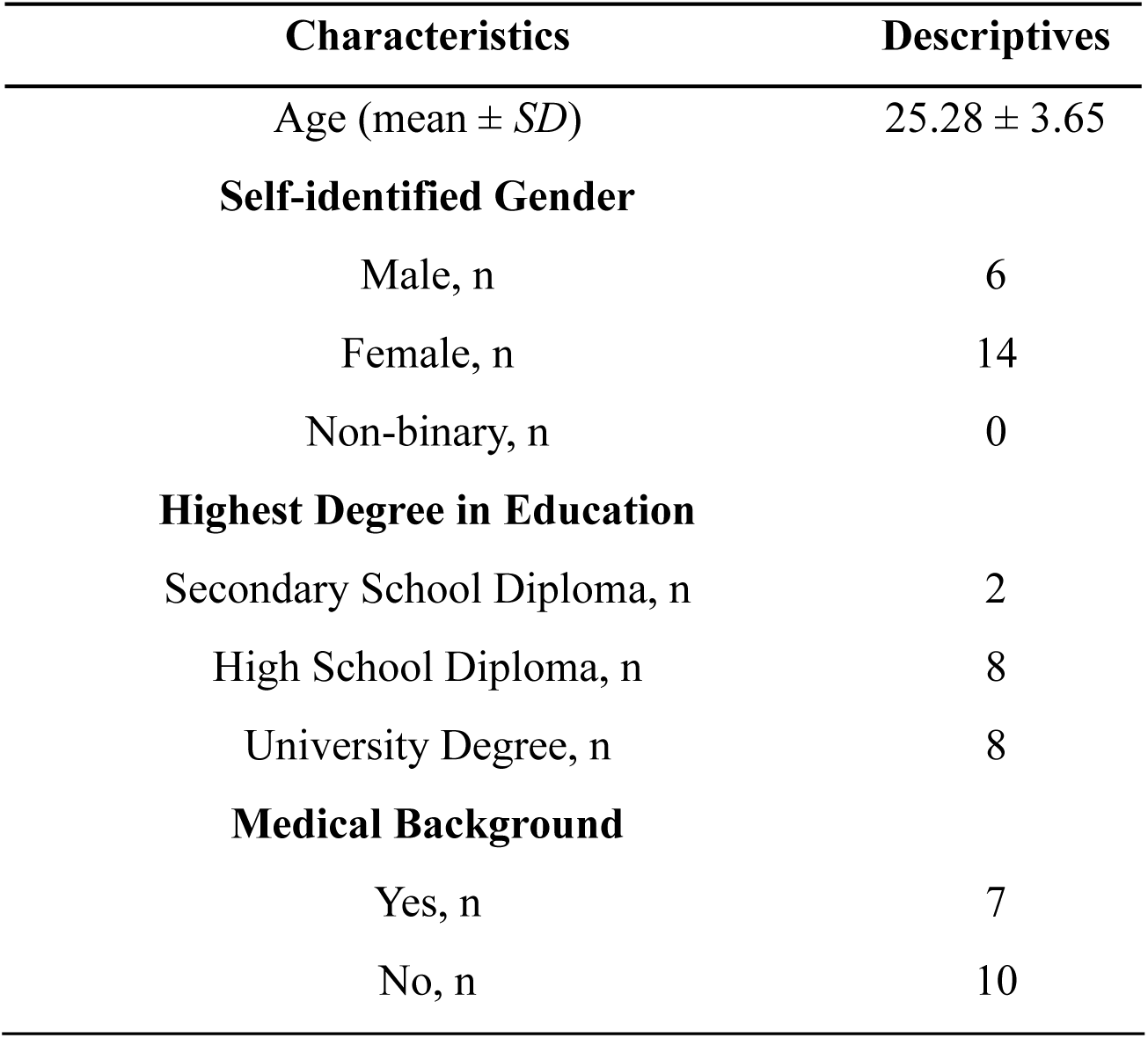
Sample characteristics Experiment 1 (n = 20). Table 1 reports the sample characteristics of the participants of Experiment 1. We report age (mean ± standard deviation), self-identified gender, education and the binary variable medical background. Participants reported a medical background when they worked or studied in medicine or related fields like e.g. nursing.

Study procedures were in line with the Declaration of Helsinki. The ethics committee of the Hamburg medical chamber approved the study. Participants that finished the study received an expense compensation of 15€ for participating in a one-hour experiment.

#### 2.2.2 Experimental design

The experiment was divided into two phases: observation phase and expression phase. In the observation phase participants were instructed to watch videoclips of a novel blood-pressure measurement approach, which was highly precise yet painful. They were told to watch the patient’s pain-expression as well as the screen, visible in the video. Participants were not instructed about the condition-design and cue/pain contingencies. The screen in the video showed a matrix of 3 horizontal grey bars (rectangles) stacked vertically (see Fig 1 b). Each bar had a colored portion (cue) of different length and position and a black slider at the left-hand side of the bar. At the start of each trial, one of three sliders (depending on the trials’ condition either P, U or N) started moving from the left-hand side of the bar to the right. In the P condition, the pain expression of the demonstrator was made predictable by the slider crossing the colored part of the bar. Here, the colored portion of the bar served as a predictive cue. In U, the slider position had no predictive value for the pain expressed by the demonstrator, i.e. the demonstrator’s pain responses were visible at random slider positions in each video clip. Therefore, the colored part of the bar served as an unpredictive cue in this condition. In the N condition, no pain expression was visible (see figure 1 b for an overview of slider/pain relation), while the slider was crossing the entire bar. Each video-clip lasted from the moment the slider started moving till it reaching its final position on the right-hand side of the grey bar (24 s). Note that each trial contained only one active condition, i.e. only one slider per trial moved. Participants were asked to rate the pain intensity for each clip. In total participants completed 12 trials in each condition.

**Figure 1:**
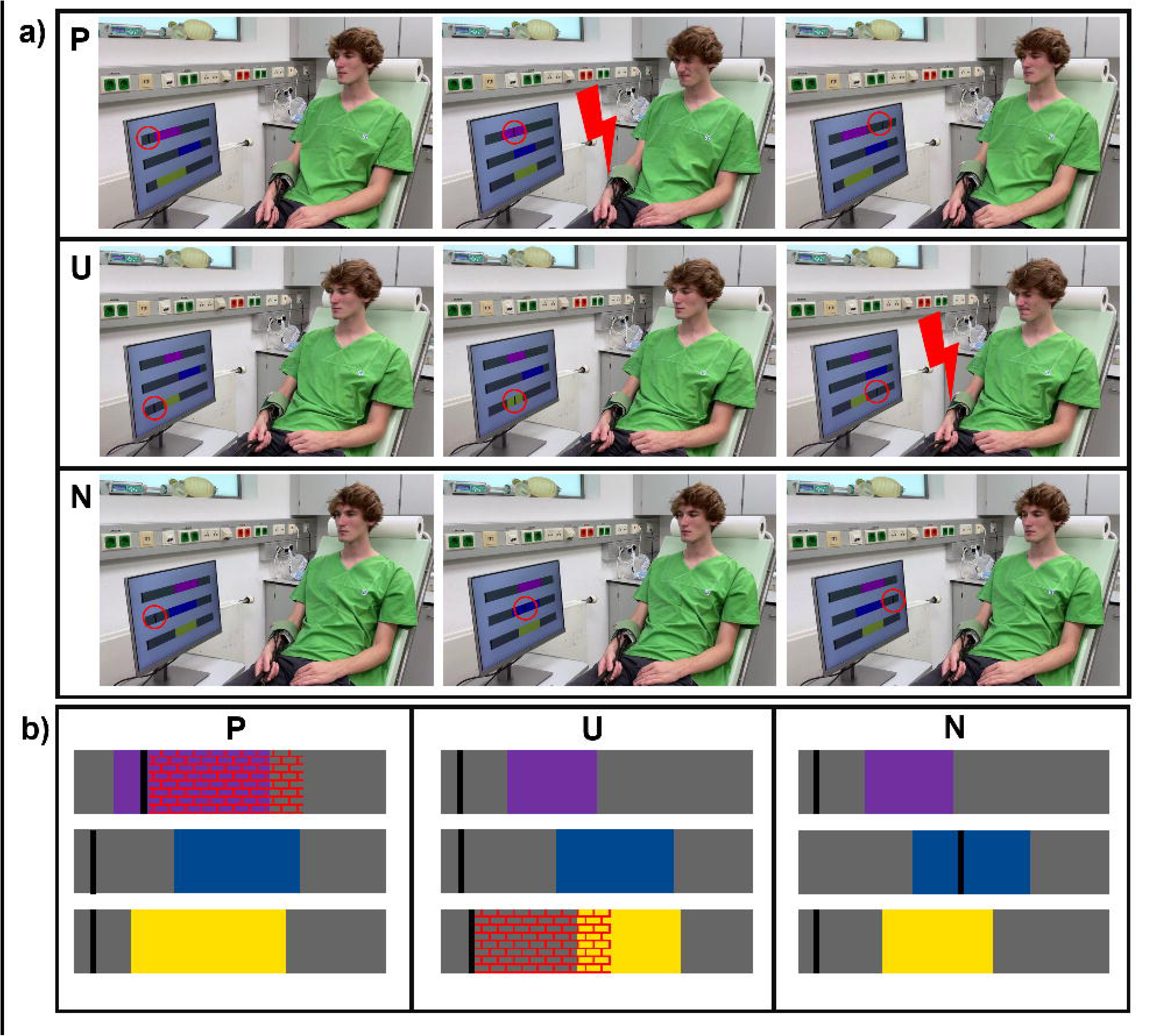
Experimental Design. a) shows three extracted frames from the P, U and N stimulus material respectively. To increase visibility, the cue position is marked with a red circle and the painful stimulation is marked with a red bolt. These markups were not visible for the participants. In P, the demonstrator is visible at rest while the slider passes the grey parts of the rectangle before and after the cue. The demonstrator is in pain during the cue (purple part of the rectangle). In U, the demonstrator is visible at rest while the slider passes the grey part of the rectangle before the cue and during cue (yellow part of the rectangle). The demonstrator is in pain during the passing of the grey part of the rectangle after the cue. In N, the demonstrator is at rest at all slider positions. b) gives an overview of the condition design. Each condition is revealed at stimulus onset, as only one slider starts moving. Unpredictable and predictable threat, thus, can be anticipated at each condition onset. The crosshatched red lines mark the typical pain position in P and a possible pain position in U. The crosshatched lines were not visible for the participants.

In the second phase, the expression phase, participants were directly confronted with what was previously visible on the demonstrator’s screen (as depicted in Fig 1 b). Similarly, the trials lasted from the moment the slider started moving till it reached its final position on the right-hand side of the grey bar (24 s). In Experiment 1, participants did not have a pain-inflicting device attached to them and were informed that no pain would be administered. Participants were asked to answer questions regarding their expectations about painful stimulation in each clip (see 2.2.3 Ratings). In total, participants completed 12 trials in each condition. The inter-trial interval included a fixation cross that was displayed for 5 seconds. Trial sequence was pseudorandomized. Three trials including one N, one P and one U condition were randomly shuffled. The shuffling process was repeated for the 12 appearances of the conditions.

#### 2.2.3 Ratings

During the observation phase, participants were asked to rate the observed pain intensity by placing a marker on a visual analog scale on a continuous line with two endpoints of no pain (1) on the left and strong pain (100) on the right. The marker was moved by pressing the arrow keys (‘left’ and ‘right’ arrow key). The rating query appeared while the video-clip was paused and blurred during the pain-expression of the demonstrator. In the N condition, the participants rated the same question while the video was paused and blurred, but no pain was visible. Following confirmation of the rating by pressing the arrow key ‘up’, the blur was removed from the video and the clip continued playing until the end. There was no time limit for response confirmation.

During the expression phase, participants were asked to rate their threat expectation and certainty by placing a marker on a visual analog scale on a continuous line with two endpoints of certainly no pain (-100) on the left and certainly pain (100) on the right. The middle point was labeled as uncertain (0). Participants rated at three time-points within a trial: condition-onset (slider position 2,5 seconds before the colored part of the bar), cue (slider position in the middle of the colored part of the bar) and condition-offset (slider position 2.5 seconds after the colored part of the bar). Note that due to a technical error in the script, the first four participants were not asked for condition-offset-expectation (condition-offset-expectation n = 16).

For each expectation-rating, participants were asked to provide two ratings: Their expectation that the demonstrator in the video would receive a painful stimulation if the demonstrator was shown the same screen currently displayed to the participant (other rating) and their expectation that they themselves would receive a painful stimulation if they were attached to the same device used on the demonstrator in the video.

#### 2.2.4 Stimuli

33 videoclips for P and U each with randomized pain-onset and -duration were recorded. The videos show a demonstrator who is watching a screen while receiving painful heat-stimulation. The demonstrator wears a cuff around his right lower forearm (see Fig 1 a) and the heat stimulation device (a thermode) is placed under the cuff. The response of the demonstrator to the heat stimulation reflected the painful experience. In order to choose video clips for the experiment comparable in pain expression, three individuals rated, independently, the presence of the following five components: lip press, head protrusion, eye closure, forehead wrinkle and masseter contraction. Furthermore, pain visibility duration (seconds) and pain-intensity (scale from 1-10, i.e. no pain resp. very strong pain) were rated. All clips that featured at least four pain components (marked as present by at least one of the three reviewers) entered a propensity-score based matching. Propensity score was calculated by the mean rated pain-intensity and duration. The caliper width of the matching was reduced until a subset of 12 clips for each condition remained for Experiment 1. The 12 included clips (Table 2 describes mean and range for pain-, cue duration and pain/cue overlap) did not differ significantly in pain-intensity (*p* = 0.690) and visible pain duration (*p* = 0.918) in Experiment 1.

**Table 2.**
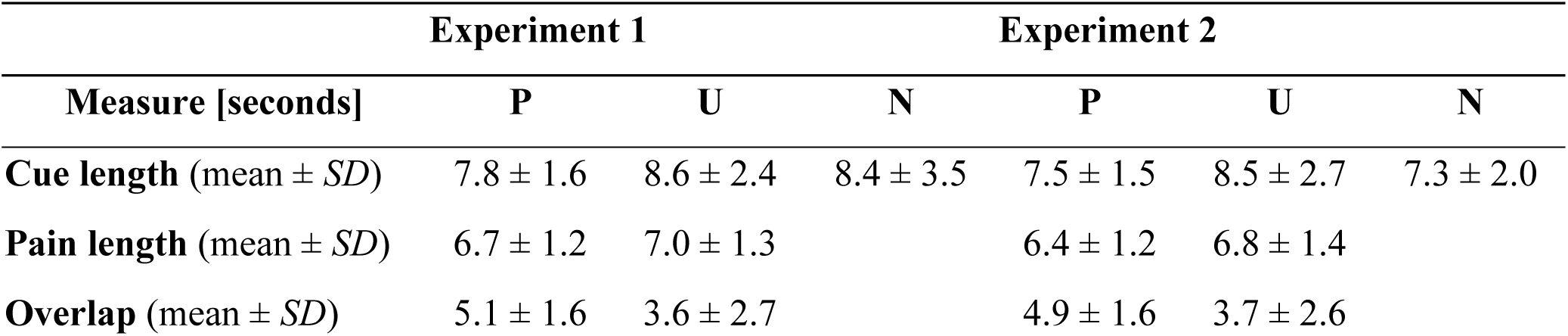
Stimuli of Experiment 1 and 2: Descriptives of measures characterizing the video stimuli. Table 2 reports a description of the video stimuli used in the experiment. We report the length of the cue, the length of the visible pain and the overlap of cue and pain. Measures are given in seconds.

### 2.3 Experiment 2

#### 2.3.1 Participants and procedure

GPower 3.1.9.7 (RRID:SCR_013726) was used to estimate the sample size for Experiment 2. Based on our findings from Experiment 1 (N=20), the relevant interaction yielded an effect size of η²_p_ = 0.331, which results in a power of >99 % for 20 participants, given an alpha rate of 0.05. Thus, we determined the sample here to be 25 participants including dropouts. 25 healthy participants (see Table 3 for descriptives) were recruited by online and black-board postings at the University Medical Center Hamburg. Eligibility criteria were the same as for Experiment 1. Additionally, participants unable to undergo fMRI scanning due to ferromagnetic or electrical implants as well as unspecified foreign bodies were excluded from study-participation.

**Table 3.** Sample characteristics Experiment 2 (n = 23). Table 3 reports the sample characteristics of the participants of Experiment 2. We report age, self-identified gender, education and the binary variable medical background. Participants reported a medical background when they worked or studied in medicine or related fields like e.g. nursing. Furthermore, we report the computed variables from the STAI and ACME questionnaires. Mean and standard deviation are reported.

| Characteristics | Descriptives |
| --- | --- |
| Age (mean ± *SD*) | 27.3 ± 6.14 |
| **Self-identified Gender** |  |
| Male, n | 13 |
| Female, n | 9 |
| Non-binary, n | 1 |
| **Highest Degree in Education** |  |
| Secondary School Diploma, n | 3 |
| High School Diploma, n | 12 |
| University Degree, n | 8 |
| **Medical Background** |  |
| Yes, n | 5 |
| No, n | 18 |
| **Questionnaires** |  |
| STAI – State (mean ± *SD*) | 33.6 ± 7.9 |
| STAI – Trait (mean ± *SD*) | 32.2 ± 7.3 |
| ACME – Cognitive Empathy (mean ± *SD*) | 27.2 ± 8.4 |
| ACME – Affective Resonance (mean ± *SD*) | 20.7 ± 8.9 |
| ACME – Affective Dissonance (mean ± *SD*) | 20.8 ± 11.4 |

Study data was collected in a single appointment of approximately 120 minutes. The experimental procedure followed a similar design to Experiment 1 with the exception that Experiment 2 was conducted in the fMRI scanner. Also, in Experiment 2 skin conductance response [SCR] was collected to assess the psychophysiological response. Furthermore, in Experiment 2 participants filled out the State-Trait Anxiety Index [STAI] (Spielberger et al., 1970) and the Affective and Cognitive Measure of Empathy [ACME] (Vachon & Lynam, 2016).

Neuroimaging acquisition consisted of two functional imaging blocks, each taking approximately 25 minutes (observation and expression, see 2.3.3 Experimental Design). Subsequently a T1 weighted image (ca. 5 minutes) for co-registration and a T2 weighted image (ca. 10 minutes) were collected. The data from the T2 weighted image is not reported in this study.

Participants who finished the study received an expense compensation of 30 € for participating in a two-hour experiment.

#### 2.3.2 Experimental design and ratings

Although the two-phase experimental design did not differ substantially from Experiment 1, there were some differences of note. In Experiment 2, participants rated pain intensity in the observation phase and threat expectations in the expression phase only in three of the twelve trials. The other nine trials ran without a rating interruption, allowing for the analysis of neural correlates. A single video clip from both P and U, matched in pain-intensity and -duration was chosen and used for each of the three rating trials.

In contrast to Experiment 1, participants in Experiment 2 wore an identical cuff, as seen on the demonstrator, and were told that it might inflict some unpleasant but not highly painful stimulation during the expression phase. However, no painful stimulation occurred during the expression phase. Furthermore, participants rated their baseline expectation, i.e. expectations towards N, P and U cues and condition-onsets prior to learning, before the observation phase.

All answers were collected by using a 4-button box. The slider-scales used in Experiment 2 were identical to the ones from Experiment 1. In the expression phase, participants were only required to rate their threat expectation and certainty regarding themselves being in the same situation as the demonstrator before. No expectations about the demonstrator were collected.

#### 2.3.3 Stimuli

For Experiment 2, caliper width was further reduced to reduce the subset to 10 video clips. The 10 included clips did not differ significantly in pain-intensity (*p =* 0.612) and visible pain duration (*p =* 0.912) in both subsets respectively (Table 2 describes mean and range for pain-, cue duration and pain/cue overlap).

#### 2.3.4 Image acquisition

All images were acquired using a 3 Tesla Siemens Prisma scanner (Siemens AG; München, Germany) at the Institute for Systems Neuroscience at the University Medical Center Hamburg-Eppendorf. For this study, echo-planar imaging [EPI] sequences sensitive to blood oxygenation level-dependent changes were acquired for the functional analysis. 52 axial slices of 1.5 mm x1.5 mm x1.5 mm voxels were acquired (TR = 1848 ms, TE = 30 ms, flipangle = 70 °). Subsequently, an anatomical image was acquired (T1 weighted 3D image, magnetization prepared rapid acquisition gradient echo image, 1 mm x 1mm x 1mm voxel size, TR = 7.1 ms, TE = 2.98 ms, TI = 1100 ms, matrix = 256 × 256, 240 slices, flip angle = 9 °).

#### 2.3.5 SCR

SCR was collected using two palmar electrodes on the participants left hand. Spike2 software (Cambridge Electronic Design Limited, Cambridge, England, RRID:SCR_000903) was used for data collection. Data was acquired with a sampling rate of 1000 Hz. MRI-pulses as well as pulses for condition-, cue- and pain-onsets were collected to synchronize the evoked responses with the experiment’s conditions.

### 2.4 Statistical analysis

#### 2.4.1 Behavioral analysis

In both Experiments (1 and 2) mean threat expectation ratings before and during cue presentation from the expression phase were analyzed in a repeated measures ANOVA with condition (P, U, N) and cue/condition-onset (cue-expectation, condition-onset-expectation) as factors. For post-hoc comparison we aimed to characterize an interaction between condition and cue vs. condition-onset. To that end, the ‘cue-prominence’ was calculated for all conditions separately by subtraction of the condition-onset-expectation from the cue-expectation. Cue prominence was compared among the conditions in paired-sample *t*-tests (P vs. N, U vs. N, P vs. U). In Experiment 1 a similar repeated measures ANOVA and post-hoc *t*-tests were calculated for expectation of pain for the demonstrator. To account for beliefs held by participants unrelated to the observable predictability of the conditions (i.e. baseline), we report pre-observation ratings for threat expectation and compare them similarly across conditions in Experiment 2. Additionally, condition-offset-expectation (slider-position behind colored part of the bar) were compared in a univariate ANOVA among the conditions (P, U, N) in both Experiment 1 and 2. Significant effects were followed up using paired-sample *t*-tests. Furthermore, mean pain ratings from the observation phase were compared among the pain-conditions in a paired-sample *t*-test (P vs. U).

Alpha-level was set to *p* < 0.05. Multiple comparisons were accounted for through Bonferroni-Holm correction. For the ANOVAs, sphericity was assessed using Mauchly’s test, and in cases of violation, Greenhouse-Geisser-corrected values were reported. All statistical analyses were conducted using JASP 0.18.3.0 (RRID:SCR_015823). Matlab 2023b (Mathworks Inc., Natick, Massachusetts, USA, RRID:SCR_001622) was used to read-out logfiles and count the participants’ answers.

#### 2.4.2 fMRI analysis

Participants were checked for excessive movement prior to analysis (preregistered exclusion criterium = 3x voxel-size lead to deletion of affected volumes, participants exceeding 25 % data quality issues were excluded from analysis). For preprocessing, EPIs were realigned to correct for motion and the six resulting movement parameters were included in the first-level analysis. Subsequently, images were coregistered to the anatomical image (reference: T1, source: EPI) and normalized into MNI space using SPM 25 defaults.

In both experimental phases of Experiment 2 (observation and expression) cue and condition-onset regressors were modeled following the focus of the behavioral analysis of cue- and condition-onset-expectation. To prevent cue-, condition- and pain-regressors from being colinear, condition-onset-regressors were modeled as event regressors at the onset of slider movement and cue regressors as event regressors in the moment the slider started the passage of the colored part of the bar. Pain-regressors were modeled as event-regressors at the onset of pain-expression, as rated by three lab members for each video clip (mean of rated onsets). For N, whose stimulus videos did not contain visible pain in the demonstrator, a no-pain event regressor was modeled starting approximately (in a random interval) 2 s after the cue-regressor to allow to compare social information processing (pain>no pain contrasts). A box-car regressor was modeled for the whole 24 s of each videoclip representing sustained condition activations through the experimental condition. ITI, instructions and ratings were modeled as separate box-car regressors. Regressors were convolved with the hemodynamic response function in the first level analysis. Temporal autocorrelations were accounted for through AR1. The resulting beta images were smoothed using a 6 mm FWHM Gaussian kernel.

For the second-level analysis a one-factor (3 levels: experimental condition) full factorial design was calculated for each regressor separately: cue, condition-onset and condition, both in the observation phase and the expression phase. Furthermore, a two-factor full-factorial design with one factor comprising cue and condition-onset and the other factor the three experimental conditions was applied to analyze neural substrates of cue-prominence. We report the following, pre-registered, *t*-contrasts for the one-factor analyses: P>N, U>N, P>U, U>P. For the cue-prominence analyses, we report a pre-registered interaction contrast with cue >condition-onset in P and cue < condition-onset in U. For the analysis of pain related activation, we report a secondary *t*-contrast P&U>N for the analysis of neural correlates of pain-expression in general. In a subsequent analysis, following Marschner and colleagues (2008), the same first-level regressors were used with linear time-modification over the course of the trials, which was again pre-registered. We report the contrast P>U (i.e. increasing activation in P and decreasing activation in U) and U>P (i.e. vv.).

For all analyses, we report activation of 3 predefined ROIs (right and left separately): amygdala, hippocampus, insular cortex from the Harvard-Oxford Cortical Structural Atlas (RRID:SCR_001476). Clusters that surpass peak-voxel FWE correction at *p* < 0.05 in small volume correction (across the three ROIs bilaterally) are reported. For amygdala clusters, ROIs of the basolateral and centromedial subregions of the amygdala [BLA and CMA respectively] were used to discern activations in mechanistically discrete amygdala subregions. Single-sided BLA and CMA masks (Amunts et al., 2005) were generated using the Anatomy toolbox (V3, RRID:SCR_013273). SPM25 (RRID:SCR_007037) was used for the preprocessing and analysis of the fMRI data.

#### 2.4.3 SCR analysis

The collected SCR data from the experiment was analyzed using PSPM7.0 (Bach & Friston, 2013). Raw data was split into two experimental phases (observational learning and direct test) and each dataset was trimmed from the onset of the first trial to the last MRI-pulse. Trimmed data was assessed visually for data quality. Participants with at least 50 % noisy- or zero-line data were excluded from further analysis. A general-linear model (GLM) for the analyses of SCRs, implemented in PSPM, was constructed. The GLM included regressors for condition-onset, cue onset (and pain onset in the observation phase). All regressors were modeled and convolved with the canonical skin conductance response function (Bach et al., 2010). Similar to the design of the rating analysis, a repeated measures ANOVA with the factors condition (3 levels) and cue/condition-onset (2 levels) was conducted. Significant factors were followed up using paired-sample *t*-tests. Pain related skin-conductance alterations were compared using paired-sample *t*-tests among the three conditions.

## 3. Results

### 3.1 Sample characteristics

In Experiment 1, the 20 recruited participants all entered the analysis. A description of the study sample can be found in Table 1.

In Experiment 2, 25 participants were recruited. One was excluded due to technical problems with stimulus presentation and erroneous data collection. A second participant was excluded, due to excessive movement during acquisition preventing the generation of a usable T1 image for co-registration.

### 3.2. Behavioral results

#### 3.2.1 Expectation ratings during the expression phase

##### 3.2.1.1 Experiment 1

First, we tested the central hypothesis that participants can learn the temporal predictability of cues for the aversive outcome by observation. To this end, we tested participants’ threat expectation for firsthand outcomes in P vs. U in the expression phase. Indeed, the repeated-measurement ANOVA of rated threat expectation revealed a significant influence of the condition (*F* = 15.175, *df* = (2, 38), *p* < .001, *η²_p_* = 0.444), cue/condition-onset (*F* = 62.033, *df* = (1, 19), *p* < .001, *η²_p_* = 0.766) and their interaction (*F* = 9.383, *df* = (2, 38), *p* < .001, *η²_p_* = 0.331) (see Table 4). Post-hoc *t*-tests to follow up on this interaction compared cue and condition-onset expectation, i.e., ‘cue-prominence’ (cue-expectation – condition-onset-expectation) between conditions revealed higher cue-related expectation in P (*t =* 3.994, *df* = 19, *p* = 0.00078, *pHolm* = 0.00234) and U (*t =* 2.449, *df* = 19, *p* = 0.024, *pHolm* = 0.048) as compared to N. The comparison of the cue-prominence between P and U revealed a trend of a higher cue-prominence in P (*t =* 1.997, *df* = 19, *p* = 0.06, *pHolm* = 0.06). Table 5 and Table 6 give an overview of the behavioral outcome measures and the statistical analyses. Plots of the behavioral results can be found in Figure 2. We conducted an additional analysis comparing cue and mean condition expectation (mean(condition-onset, condition-offset)) using the same analytic approach, revealing similar results of higher cue prominence in P and U, compared to N, as well as higher in P compared to U (see supplementary table S1-S3). Additionally, we asked for participants’ expectation about the pain the demonstrator might receive during the expression phase, which did not match the learned threat contingencies from the observation phase. You can find this additional analysis in the supplements (supplementary text S1).

**Figure 2:**
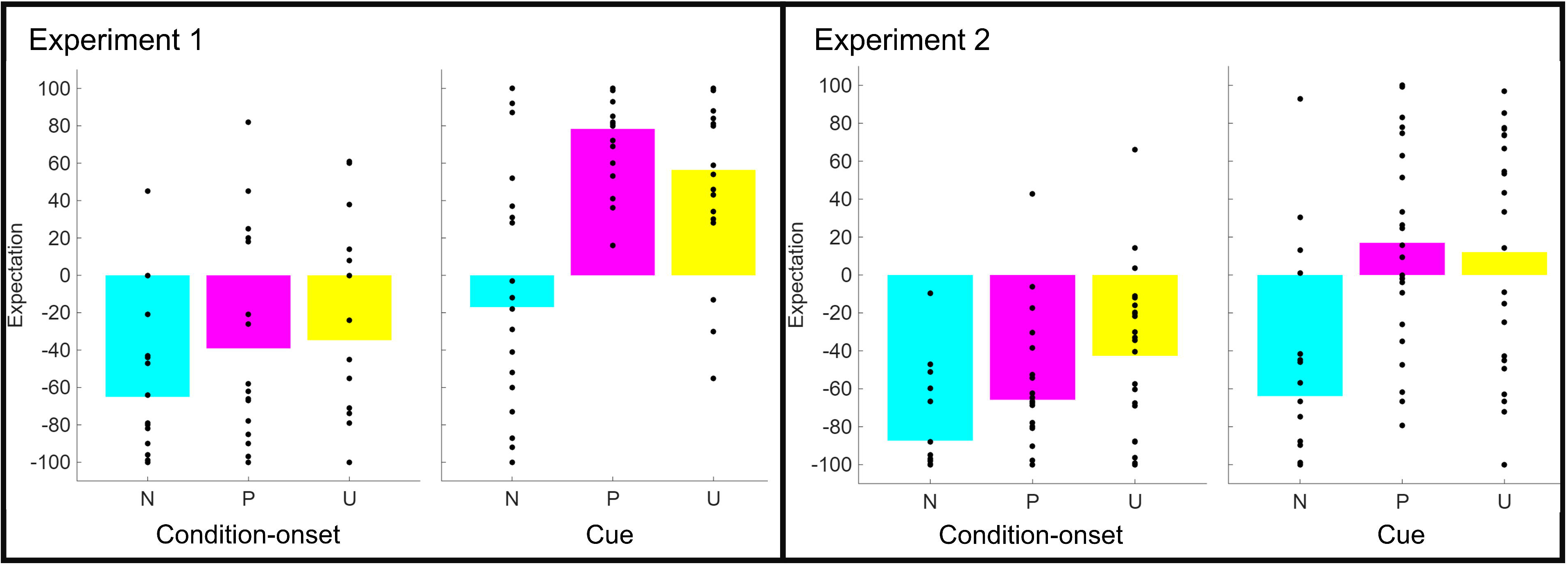
Behavioral results. Figure 2 depicts bar plots with superimposed scatter element. Bars represent the mean expectation of pain during context-onset and during cue (see 2.2.3).

**Table 4.**
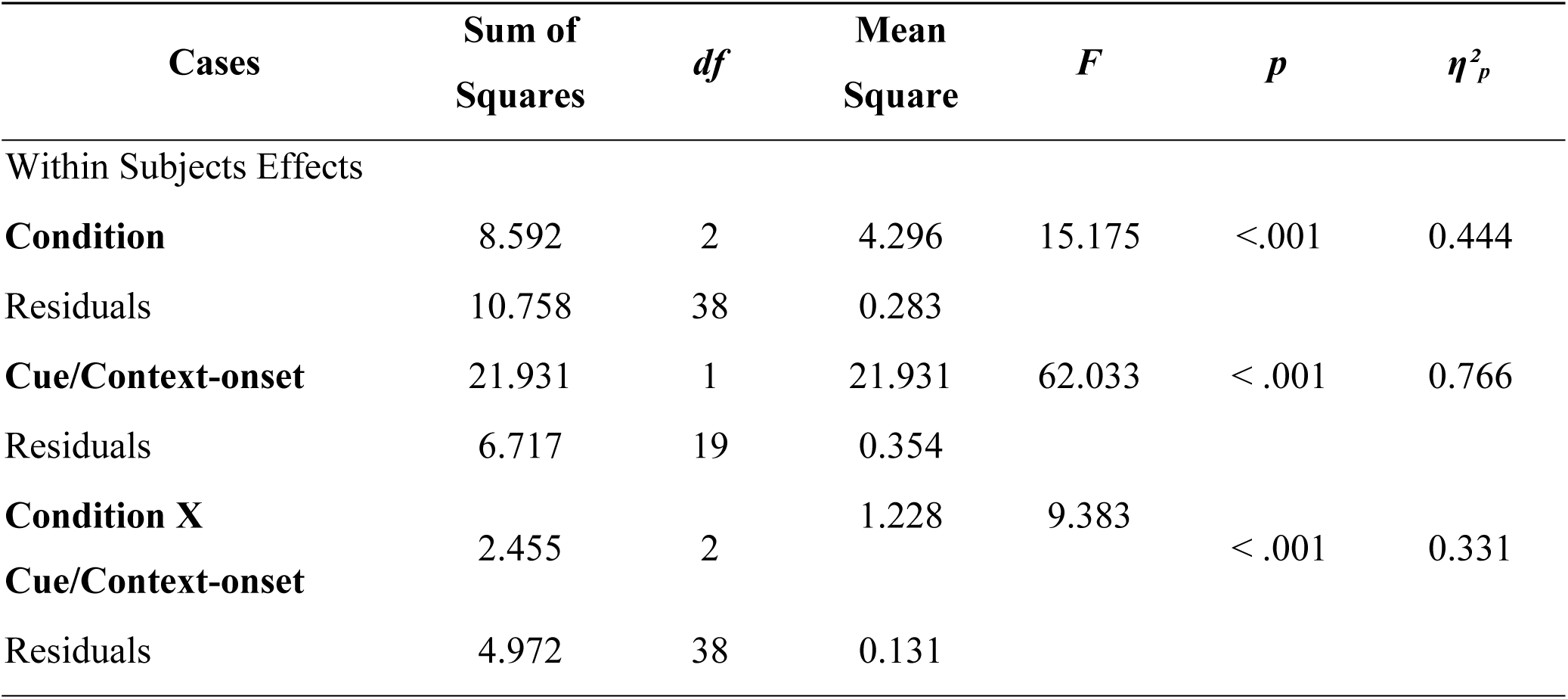
Experiment 1: Main effect repeated measures ANOVA. Threat Expectation. Table 4 reports the detailed results of the behavioral effect (threat expectation) rmANOVA (two factors: Condition: P, U, N; Timepoint: cue, context-onset) of Experiment 1. We report sum of squares, degrees of freedom, mean square, *F*-values, *p* and an estimation of the effect sizes (partial η²) for the main effects of condition and timepoint and the interaction effect.

| Cases | Sum of Squares | *df* | Mean Square | *F* | *p* | $\eta^2_p$ |
| --- | --- | --- | --- | --- | --- | --- |
| Within Subjects Effects |  |  |  |  |  |  |
| **Condition** | 8.592 | 2 | 4.296 | 15.175 | <.001 | 0.444 |
| Residuals | 10.758 | 38 | 0.283 |  |  |  |
| **Cue/Context-onset** | 21.931 | 1 | 21.931 | 62.033 | < .001 | 0.766 |
| Residuals | 6.717 | 19 | 0.354 |  |  |  |
| **Condition X Cue/Context-onset** | 2.455 | 2 | 1.228 | 9.383 | < .001 | 0.331 |
| Residuals | 4.972 | 38 | 0.131 |  |  |  |

**Table 5.**
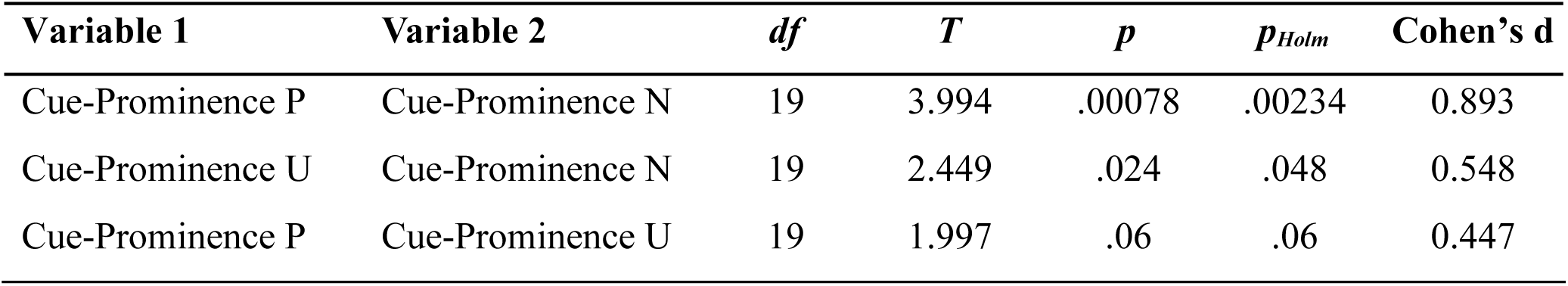
Experiment 1: Paired-sample *t*-tests. Learned cue-prominence after observation phase. Table 5 reports the posthoc paired-sample *t*-tests conducted to analyze the significant interaction effect of the rmANOVA in Experiment 1. We compare the learned cue-prominence between all three conditions. Degrees of freedom, T, *p*, *p* corrected for multiple comparisons (Bonferroni-Holm) and an estimation of effect size (Cohen’s d) are reported.

**Table 6.**
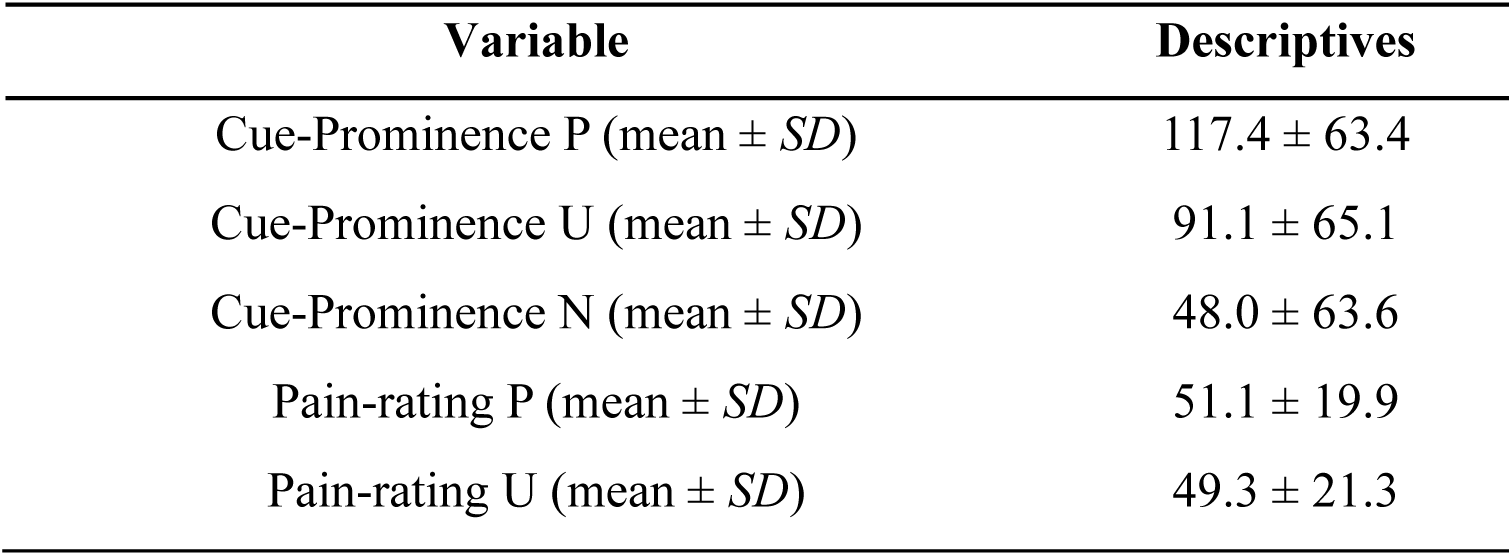
Experiment 1: Behavioral outcome measures. Table 6 reports the descriptive results of Experiment 1. We report the calculated cue-prominence for each condition (P, U, N) and the mean pain-ratings for P and U.

| Variable | Descriptives |
| --- | --- |
| Cue-Prominence P (mean ± *SD*) | 117.4 ± 63.4 |
| Cue-Prominence U (mean ± *SD*) | 91.1 ± 65.1 |
| Cue-Prominence N (mean ± *SD*) | 48.0 ± 63.6 |
| Pain-rating P (mean ± *SD*) | 51.1 ± 19.9 |
| Pain-rating U (mean ± *SD*) | 49.3 ± 21.3 |

To test if threat expectations were sustained, we compared threat expectation after the slider passed the colored cue across the conditions, which, however, only revealed a trend-wise effect (*F* = 2.772, *df* = (2, 38), *p* = 0.079, *η²_p_* = 0.156, univariate ANOVA, one factor: condition). Nevertheless, participants’ descriptive ratings matched the expected pattern, with lowest threat expectation for N (mean ± *SD*: -82 ± 31), a medium threat expectation for P (mean ± *SD*: -66 ± 30) and the highest, sustained expectation of an aversive outcome in U (mean ± *SD*: -53 ± 43).

##### 3.2.1.2 Experiment 2

In Experiment 2 we were able to reproduce and further advance the results from Experiment 1. Similar to Experiment 1, the repeated measures ANOVA of threat expectation in the expression phase revealed a significant main effect of condition (*F* = 22.642, *df* = (2, 44), *p* < .001, *η²_p_* = 0.507), cue/condition-onset (*F* = 32.291, *df* = (1, 22), *p* < .001, *η²_p_* = 0.595) and their interaction (*F* = 13.158, *df* = (2, 44), *p* < .001, *η²_p_* = 0.374) (see Table 7 for Greenhouse-Geisser corrected values). Pre-registered post-hoc comparisons again revealed that the cue-prominence was increased in P (*t =* 4.443, *df* = 22, *p* = 0.0002, *pHolm* = 0.0006) and U (*t =* 2.814, *df* = 22, *p* = 0.01, *pHolm* = 0.02) in comparison to N and we found an increased cue-prominence in P as compared to U (*t =* 2.808, *df* = 22, *p* = 0.01, *pHolm* = 0.01) (see Table 8, see figure 2). We found comparable threat expectation for the cue in P and U, but an increased threat expectation at condition-onset in U as compared to P (Cue-Expectation P vs. U: *t* = 0.891, *df* = 22, *p* = 0.382, *pHolm* = 0.382; Condition-onset-Expectation = U vs. P: *t* = 2.615, *df* = 22, *p* = 0.016, *pHolm* = 0.032). The negative rating values at unpredictable condition-onset indicate that participant’s expectation to receive pain at this timepoint is low, however, this expectation rating increases over the trial. In addition, the results also suggest that participants display uncertainty at condition-onset (ratings range includes 0).

**Table 7.** Experiment 2: Main effect repeated measures ANOVA. Threat Expectation. Table 7 reports the detailed results of the behavioral effect (threat expectation) rmANOVA (two factors: Condition: P, U, N; Timepoint: cue, context-onset) of Experiment 2. We report sum of squares, degrees of freedom, mean square, *F*-values, *p* and an estimation of the effect sizes (partial η²). ᵃ points at the violation of the assumption of sphericity (Mauchly’s test, *p* < .05), in which case Greenhouse-Geisser corrected values are reported.

| Cases | Sum of Squares | *df* | *Mean Square* | *F* | *p* | $\eta^2_p$ |
| --- | --- | --- | --- | --- | --- | --- |
| Within Subjects Effects |  |  |  |  |  |  |
| **Condition** | 9.729<sup>a</sup> | 2<sup>a</sup> | 4.865<sup>a</sup> | 22.642<sup>a</sup> | <.001<sup>a</sup> | 0.507 |
| Greenhouse-Geisser | 9.729 | 1.325 | 7.343 | 22.642 | <.001 | 0.507 |
| Residuals | 9.453 | 44 | 0.215 |  |  |  |
| Greenhouse-Geisser | 9.453 | 29.148 | 0.324 |  |  |  |
| **Cue/Context-onset** | 9.926 | 1 | 9.926 | 32.291 | < .001 | 0.595 |
| Residuals | 6.762 | 22 | 0.307 |  |  |  |
| **Condition X Cue/Pr-Cue Context** | 2.022 | 2 | 1.011 | 13.158 | < .001 | 0.374 |
| Greenhouse-Geisser | 2.022 | 1.786 | 1.132 | 13.158 | <.001 | 0.374 |
| Residuals | 3.381 | 44 | 0.077 |  |  |  |
| Greenhouse-Geisser | 3.381 | 39.296 | 0.086 |  |  |  |

**Table 8.** Experiment 2: Behavioral outcome measures. Table 8 reports the descriptive results of Experiment 1. We report the calculated cue-prominence for each condition (P, U, N) before (initial cue-prominence) and after learning and the mean pain-ratings for P and U.

| Variable | Descriptives |
| --- | --- |
| Initial Cue-Prominence P (mean ± *SD*) | 55.7 ± 79.6 |
| Initial Cue-Prominence U (mean ± *SD*) | 31.6 ± 55.9 |
| Initial Cue-Prominence N (mean ± *SD*) | 48.2 ± 62.6 |
| Cue-Prominence P (mean ± *SD*) | 82.8 ± 60.8 |
| Cue-Prominence U (mean ± *SD*) | 54.7 ± 56.8 |
| Cue-Prominence N (mean ± *SD*) | 23.5 ± 47.9 |
| Pain-rating P (mean ± *SD*) | 62.6 ± 20.2 |
| Pain-rating U (mean ± *SD*) | 66.5 ± 21.9 |

As expected, baseline expectations before observational threat learning were not different among the three conditions (see Table 9). Similar to Experiment 1, we conducted an additional analysis comparing cue and mean condition expectation (mean(condition-onset, condition-offset), see supplementary tables S4-S6).

**Table 9.** Experiment 2: Paired-sample *t*-tests. Learned cue-prominence after observation phase. Table 9 reports the posthoc paired-sample *t*-tests conducted to analyze the significant interaction effect of the rmANOVA in Experiment 2. We compare the learned cue-prominence between all three conditions. Degrees of freedom, T, *p*, *p* corrected for multiple comparisons (Bonferroni-Holm) and an estimation of effect size (Cohen’s d) are reported.

| Variable 1 | Variable 2 | *df* | *T* | *p* | *p<sub>Holm</sub>* | Cohen's *d* |
| --- | --- | --- | --- | --- | --- | --- |
| Learned cue-prominence P | Learned cue-prominence N | 22 | 4.443 | .0002 | .0006 | .926 |
| Learned cue-prominence U | Learned cue-prominence N | 22 | 2.814 | .01 | .02 | .587 |
| Learned cue-prominence P | Learned cue-prominence U | 22 | 2.808 | .01 | .01 | .586 |

Participants sustained a differential threat expectation across conditions after the slider passed the colored cue (*F* = 5.989, *df* = (2, 44), *p* = 0.005, *η²_p_* = 0.214)). Similar to the pattern found in Experiment 1, we found an increased rating for both P (*t =* 3.941, *df* = 22, *p* = 0.00066, *pHolm* = 0.002) and U (*t =* 2.581, *df* = 22, *p* = 0.017, *pHolm* = 0.034) compared to N, however, no difference was observed between P and U (*t =* 0.313, *df* = 22, *p* = 0.758, *pHolm* = 0.758), which may indicate that the predictable condition was evaluated as potentially dangerous, even when the predictable danger signal passed.

#### 3.2.2 Pain ratings during the observation phase

In Experiment 1, participants rated a higher pain-intensity for the demonstrator in P as compared to U (see Table 6). This descriptive difference, however, did not reach statistical significance (*t =* 1.910 *df* = 19, *p* = 0.071).

In Experiment 2, our pre-registered, directed analysis revealed no difference in ratings for the demonstrator’s inferred pain between P and U (*t =* 1.384, *df* = 22, *p* = 0.180). Table 10 gives an overview of the behavioral measures taken in Experiment 2 descriptively.

**Table 10.** Experiment 2: Paired-sample *t*-tests. Initial cue-prominence before observation phase. Table 10 reports the paired-sample *t*-tests conducted to analyze the cue-prominence previous to the observational learning phase in Experiment 2. We compare the learned cue-prominence between all three conditions. Degrees of freedom, T, *p* and *p* corrected for multiple comparisons (Bonferroni-Holm) are reported.

| Variable 1 | Variable 2 | *df* | *T* | *p* | *p<sub>Holm</sub>* |
| --- | --- | --- | --- | --- | --- |
| Initial cue-prominence P | Initial cue-prominence N | 22 | 0.943 | .356 | .356 |
| Initial cue-prominence U | Initial cue-prominence N | 22 | -2.443 | .023 | .069 |
| Initial cue-prominence P | Initial cue-prominence U | 22 | 2.066 | .052 | .104 |

### 3.3 fMRI results

#### 3.3.1 Observation phase

##### Resolving cues and conditions during observational threat learning

In line with meta-analytic evidence (Jauniaux et al., 2019; Lamm et al., 2011) we found that the observation of pain across P and U elicited activation in the left insula (*t* = 5.93, *df* = [1,66], MNI = [-36, 18.5, 0.5], *p(FWE)* = 0.00006) and amygdala (*t* = 4.54, *df* = [1,66], MNI = [-28.5, 0.5, -22], *p(FWE)* = 0.0031; see Figure 3 a) – f)), when compared to observed no pain. Contrasted separately against N, observation of pain in P revealed a cluster in the left amygdala (*t* = 4.61, *df* = [1,66], MNI = [-28.5, 0.5, -22], *p(FWE)* = 0.0025) and bilateral insula (left: *t* = 5.81, *df* = [1,66], MNI = [-36, 18.5, 0.5], *p(FWE)* = 0.00009; right: *t* = 4.35, *df* = [1,66], MNI = [31.5 24.5 -2.5], *p(FWE)* = 0.0124), while observation of pain in U revealed a cluster in the left insula (*t* = 4.50, *df* = [1,66], MNI = [-36, 18.5, 0.5], *p(FWE)* = 0.0029), only. Interestingly, observation of predictable pain (P>U) was further encompassed by higher parameter estimates in the right hippocampus (*t* = 4.27, *df* = [1,66], MNI = [33, -32.5, -10], *p(FWE)* = 0.0145), when compared to observed pain that was unpredictable (see Figure 3 g) – i)).

**Figure 3:**
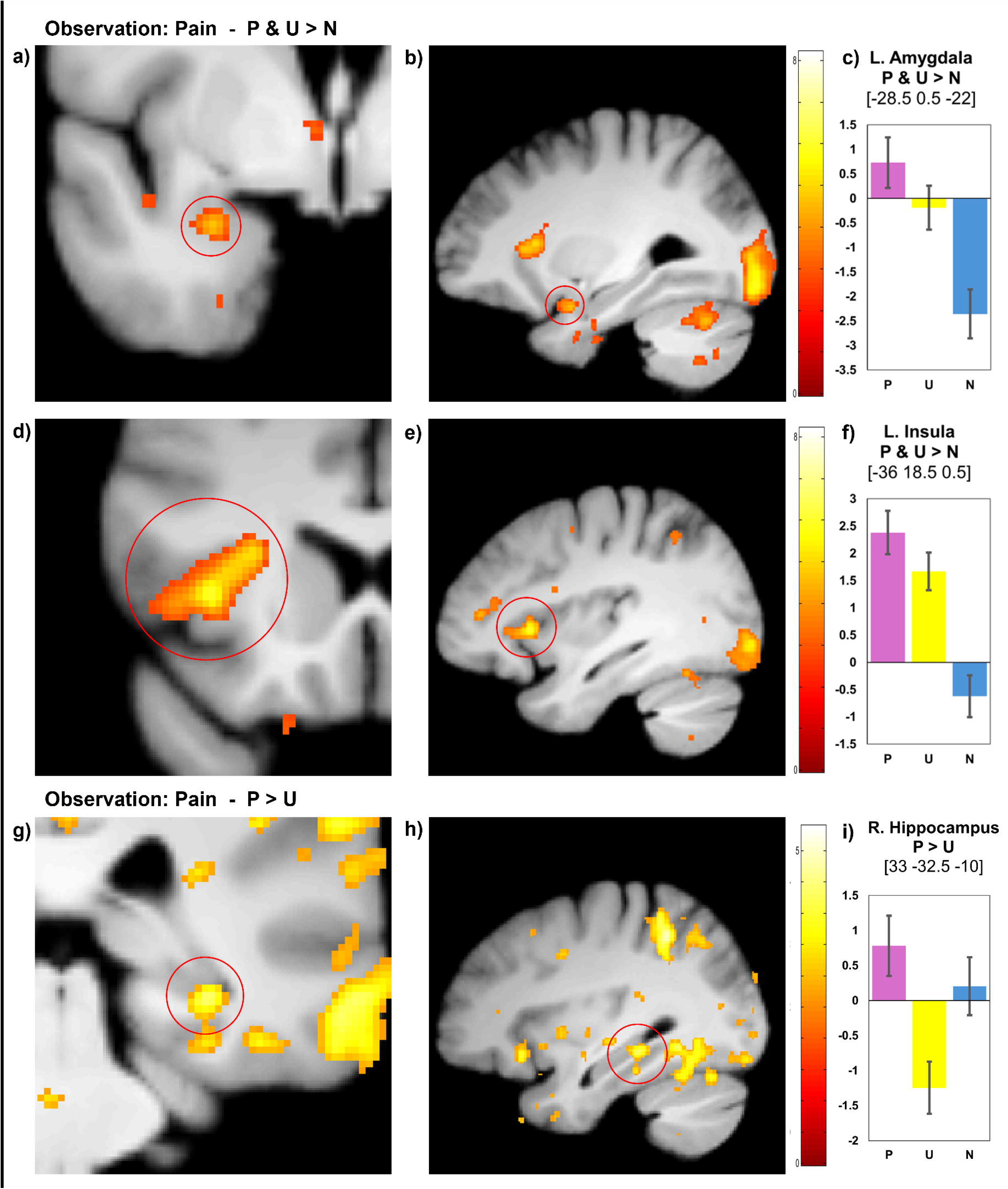
Observation phase Pain. a)-f) depict the contrast P & U > N. a) shows a 80×80 voxel zoom in coronary view of the left basolateral amygdala cluster. b) depicts a full volume sagittal view of the same cluster. c) shows diagrams of the peak-voxel ß-estimates of the left amygdala cluster. The respective cluster is highlighted with a red circle in a) and b). d) shows a 80×80 voxel zoom in coronary view of the left insula cluster. e) depicts a full volume sagittal view of the same cluster. f) shows diagrams of the peak-voxel ß-estimates of the left insula cluster. The respective cluster is highlighted with a red circle in d) and e). g) – i) depict the contrast P > U. g) shows a 80×80 voxel zoom in coronary view of the right hippocampus cluster. g) depicts a full volume sagittal view of the same cluster. i) shows diagrams of the peak-voxel ß-estimates of the right hippocampus cluster. The respective cluster is highlighted with a red circle in g) and h).

Next, we tested for neural activation that may reflect the participant’s ability to distinguish between cue and condition-onset in P vs U. Specifically, we contrasted (pre-registered) hemodynamic responses reflecting cue prominence between P and U ((PCue > PCondition-onset)>(UCue>UCondition-onset)), that mirror higher cue prominence for P, as compared to U as rated in the expression phase. This contrast only revealed a cluster trending towards FWE-corrected significance in the left (anterior) hippocampus (*t* = 3.66, *df* = [1,132], MNI = [-21, -20.5, -20.5], *p(FWE)* = 0.056) (see Figure 4 a) – c)).

**Figure 4:**
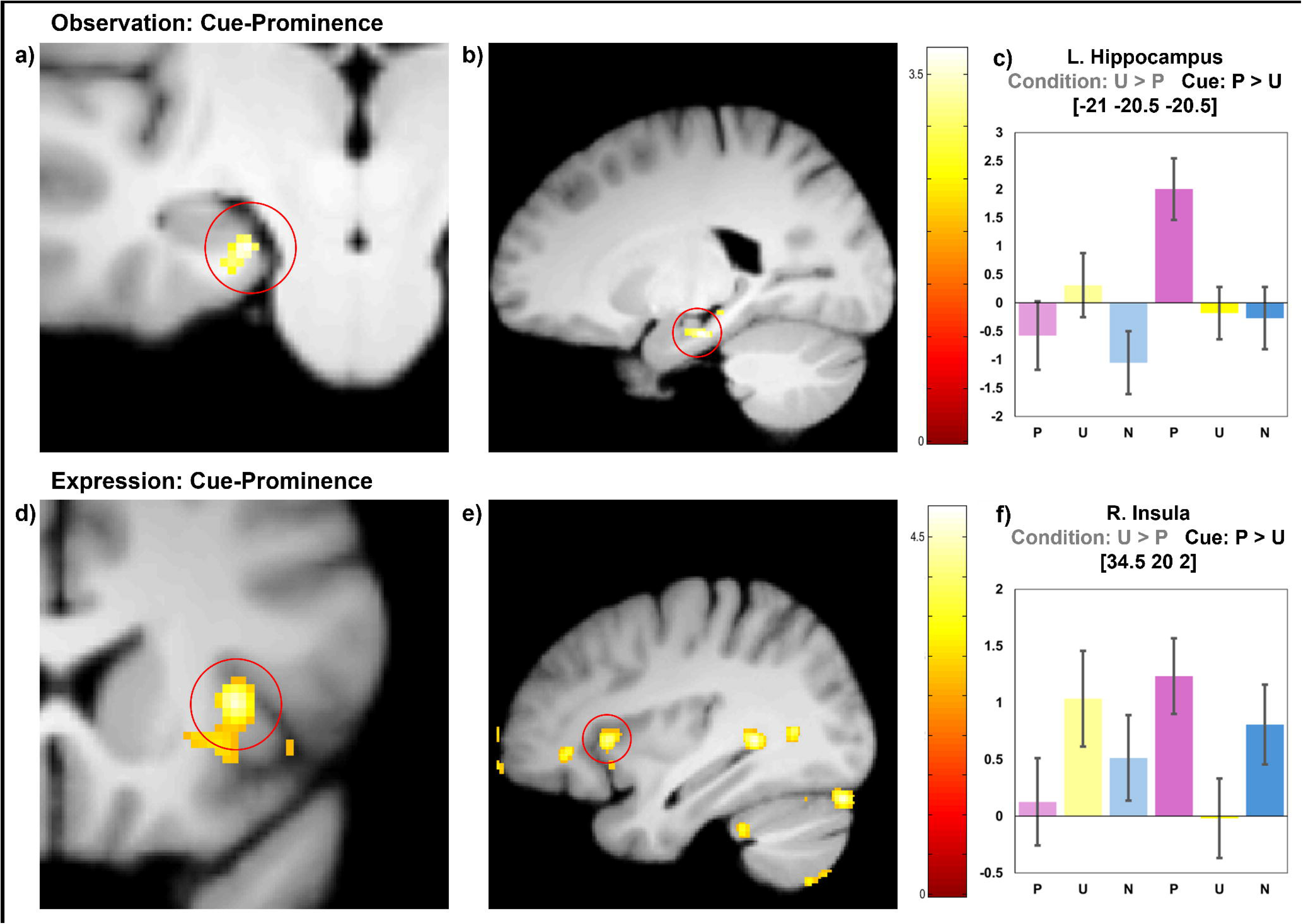
Cue Prominence. a)-c) depict the cue-prominence contrast (context: U > P, cue: P > U) of the observation phase. Note that this cluster only trends significance in FWE correction (*p* = 0.056). a) shows a 80×80 voxel zoom in coronary view of the left hippocampus cluster. b) depicts a full volume sagittal view of the same cluster. c) shows diagrams of the peak-voxel ß-estimates of the left hippocampus cluster. The respective cluster is highlighted with a red circle in a) and b). d) – e) depict the cue-prominence contrast (context: U > P, cue: P > U) of the expression phase. d) shows a 80×80 voxel zoom in coronary view of the right insula cluster. e) depicts a full volume sagittal view of the same cluster. f) shows diagrams of the peak-voxel ß-estimates of the right insula cluster. The respective cluster is highlighted with a red circle in d) and e). Note: light shaded bars in both plots of the beta estimates c) and f) represent the estimates for the respective context-onsets, dark shaded bars represent the estimates for the respective cue.

To account for learning-dependent changes during observational learning, we modeled linear changes in activity across trials. This approach was preregistered and has previously been demonstrated in both firsthand cue and context learning (Marschner et al., 2008). The interaction contrasts reflecting rated cue-prominence (contrast: linear decrease cue: P > U; linear decrease condition-onset U> P) revealed activation in the right insula (*t* = 3.76, *df* = [1,132], MNI = [45, 6.5, -7], *p(FWE)* = 0.045). This activation may reflect decreasing responses across learning to valid threat predictors, i.e. predictive cues and unpredictable condition-onsets (see Marschner et al., 2008), while activity to unpredictive cues and predictable condition-onsets was sustained or increased (see Figure 7 a) – c)). A similar pattern was found in the amygdala, specifically the bilateral CMA (*t* = 4.21, *df* = [1,66], MNI = [-18, -7, -13], *p(FWE)* = 0.009; *t* = 3.97, *df* = [1,66], MNI = [21, -5.5, -13], *p(FWE)* = 0.018) (see Figure 8 a) – c)). Here, activation decreased linearly across trials during observational learning of predictive cues. In contrast, activity remained sustained during learning of unpredictive threat cues (contrast: linear decrease P > U), which may reflect an early integration of socially derived threat information in the predictable condition.

##### Cue-specific responses during observational threat learning

To examine only cue-related activation, we contrasted activation to predictive cues of observed pain, with activation to neutral cues (P > N), which revealed activation in the posterior part of the left hippocampus (*t* = 3.90, *df* = [1,66], MNI = [-13.5, -38.5, 2], *p(FWE)* = 0.041; see Figure 5). No differences in activation in our ROIs emerged in the comparison of predictive and unpredictive cues (P > U), similar to the comparable pattern in the reports of threat expectation.

**Figure 5:**
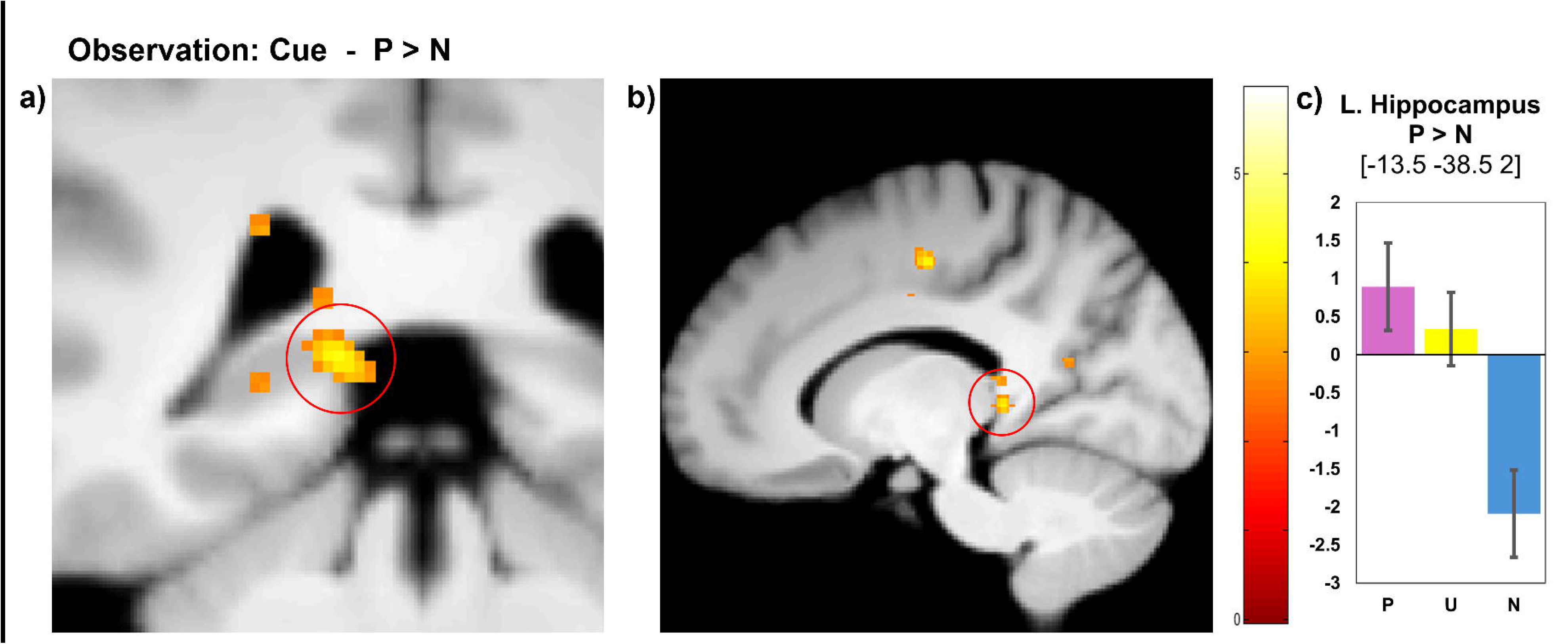
Observation phase: Cue. a)-c) depict the contrast P > N of the cue. a) shows a 80×80 voxel zoom in coronary view of the left hippocampus cluster. b) depicts a full volume sagittal view of the same cluster. c) shows diagrams of the peak-voxel ß-estimates of the left hippocampus cluster. The respective cluster is highlighted with a red circle in a) and b).

##### Condition-specific responses during observational threat learning

In the unpredictable condition, we found condition-onset related activation in the left anterior insula (*t* = 4.51, *df* = [1,66], MNI = [-36, 11, -14.5], *p(FWE)* = 0.007) when compared to the neutral condition (U > N) (see Figure 6 a) -c)). We found no differences in activation in the ROIs at condition-onset between U and P. However, the same contrast revealed sustained activation across the whole observational trial in right BLA (*t* = 3.77, *df* = [1,66], MNI = [21, -4, -20.5], *p(FWE)* = 0.027; *t* = 3.69, *df* = [1,66], MNI = [30, -4, -16], *p(FWE)* = 0.034) during the unpredictable relative to the predictable condition (U > P) (see Figure 3 d) -f)).

**Figure 6:**
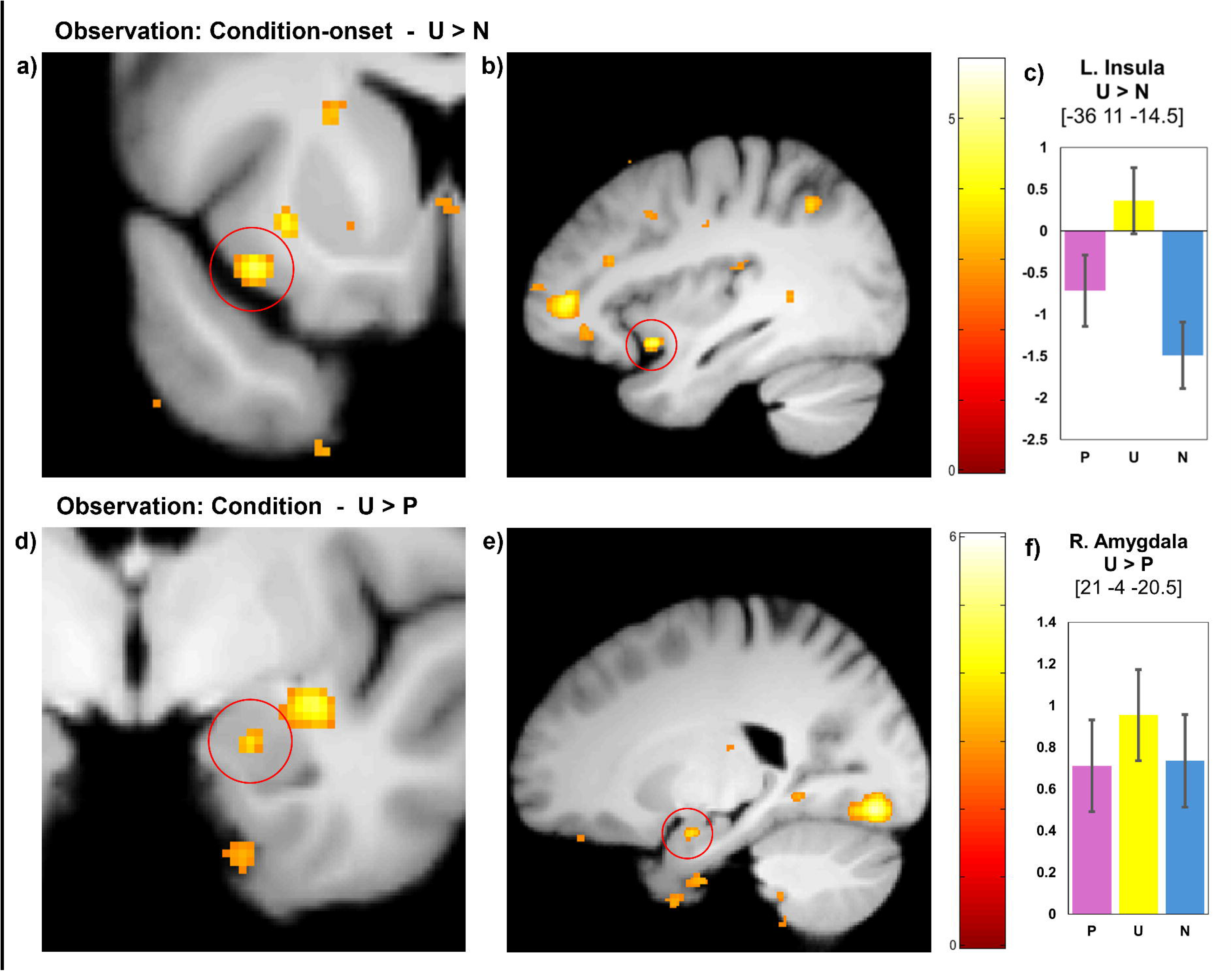
Observation phase: Context. a)-c) depict the contrast U > N of the context-onset. a) shows a 80×80 voxel zoom in coronary view of the left insula cluster. b) depicts a full volume sagittal view of the same cluster. c) shows diagrams of the peak-voxel ß-estimates of the left insula cluster. The respective cluster is highlighted with a red circle in a) and b). d) – e) depict the contrast U > P of the sustained context activity. d) shows a 80×80 voxel zoom in coronary view of the right basolateral amygdala cluster. e) depicts a full volume sagittal view of the same cluster. f) shows diagrams of the peak-voxel ß-estimates of the right amygdala cluster. The respective cluster is highlighted with a red circle in d) and e).

#### 3.3.2 Expression phase

##### Resolving cues and conditions during threat expression

The neural activation during the expression phase was hypothesized to reflect participant’s ratings of socially shaped threat expectation. Specifically, brain activity was expected to be higher for the cue relative to the condition in the predictable compared to the unpredictable condition (i.e., cue-prominence). This preregistered interaction contrast (P: cue > condition-onset, U: condition-onset > cue) revealed a significant cluster in the right insula (*t* = 4.23, *df* = [1,132], MNI = [34.5, 20, 2], *p(FWE)* = 0.009) (see Figure 4 d) – f)) and suggests a neuronal activation pattern reflecting participants’ threat expectations during the resolution of temporal threat uncertainty.

To account for changes across trials in the direct expression phase (i.e., no firsthand outcomes), we modeled linearly changing activity, as preregistered. This analysis revealed activity in amygdala subregions, the CMA and BLA that diverged between predictive cues and unpredictable condition-onsets (*t* = 4.56, *df* = [1,132], MNI = [24, -8.5, -14.5], *p(FWE)* = 0.00013), as specified in the cue-prominence interaction contrast (contrast: linear decrease cue: P > U; linear decrease condition-onset U> P). Again, activation decreased to predictive cues (compared to predictable condition-onsets) and unpredictable condition-onsets (compared to unpredictive cues) (see Figure 7 d) – f)).

**Figure 7:**
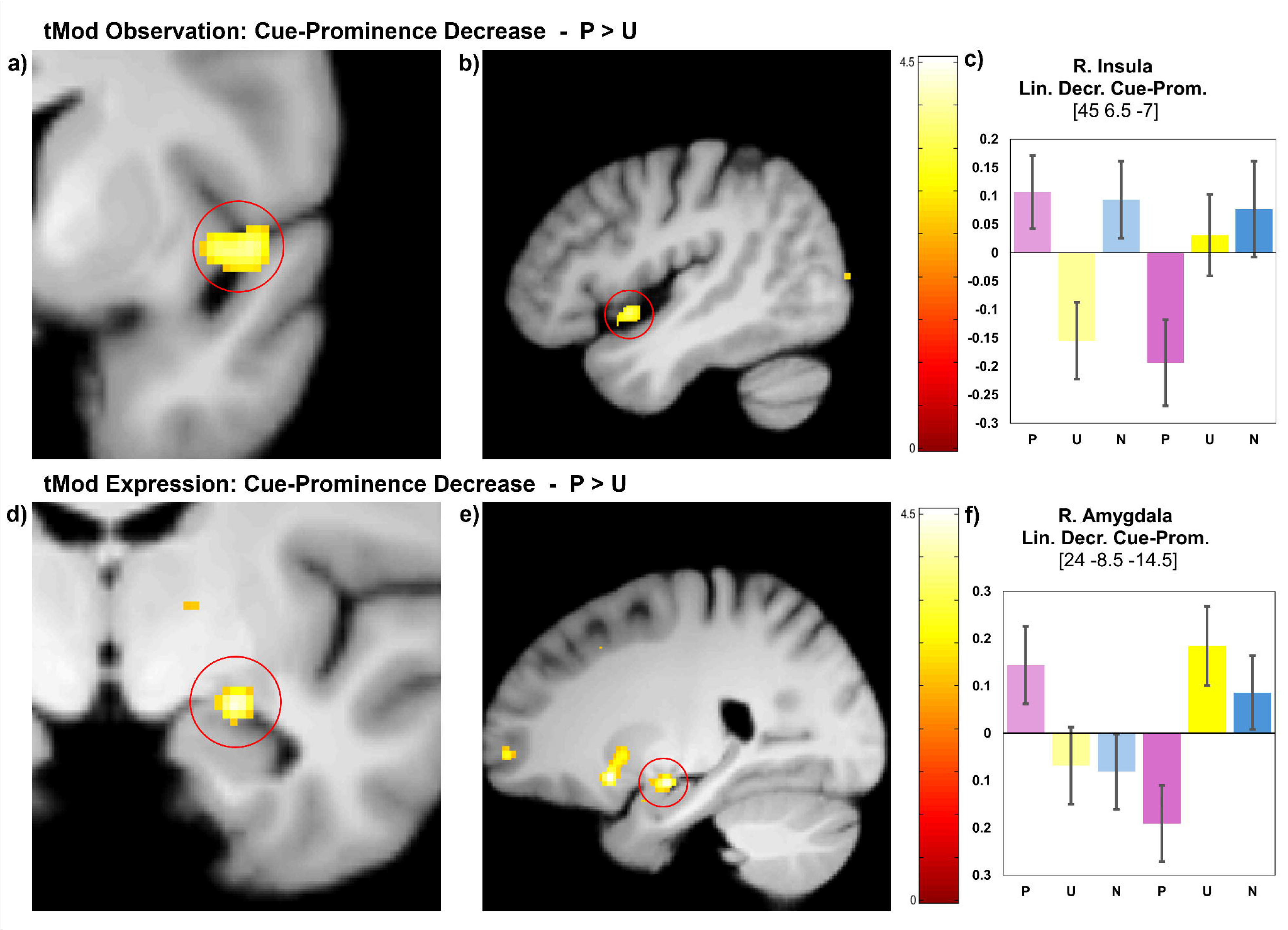
tMod cue-prominence. a)-c) depict the linearly decreasing interaction contrast of the cue-prominence in the observation phase. a) shows a 80×80 voxel zoom in coronary view of the right insula cluster. b) depicts a full volume sagittal view of the same cluster. c) shows diagrams of the peak-voxel ß-estimates of the right insula cluster. The respective cluster is highlighted with a red circle in a) and b). d)-e) depict the linearly decreasing interaction contrast of the cue-prominence in the expression phase. d) shows a 80×80 voxel zoom in coronary view of the right centromedial and basolateral amygdala cluster. e) depicts a full volume sagittal view of the same cluster. f) shows diagrams of the peak-voxel ß-estimates of the right amygdala cluster. The respective cluster is highlighted with a red circle in d) and e). Note: light shaded bars in both plots of the beta estimates c) and f) represent the estimates for the respective context-onsets, dark shaded bars represent the estimates for the respective cue.

##### Cue-specific responses during threat expression

In line with the linearly changing neural responses found in the cue-prominence contrast, we found transient CMA (*t* = 3.75, *df* = [1,66], MNI = [25.5, -8.5, -13], *p(FWE)* = 0.029) activation to predictive cues in the expression phase (see Figure 8 d) – f)). In our preregistered analysis, we found no activation in the predefined ROIs when comparing the mean responses to predictive threat-cues as compared to unpredictive cues. The comparison between unpredictive cues to predictive cues (U > P) revealed activation in the left hippocampus (*t* = 5.39, *df* = [1,66], MNI = [-19.5, -10, -23.5], *p(FWE)* = 0.0004).

**Figure 8:**
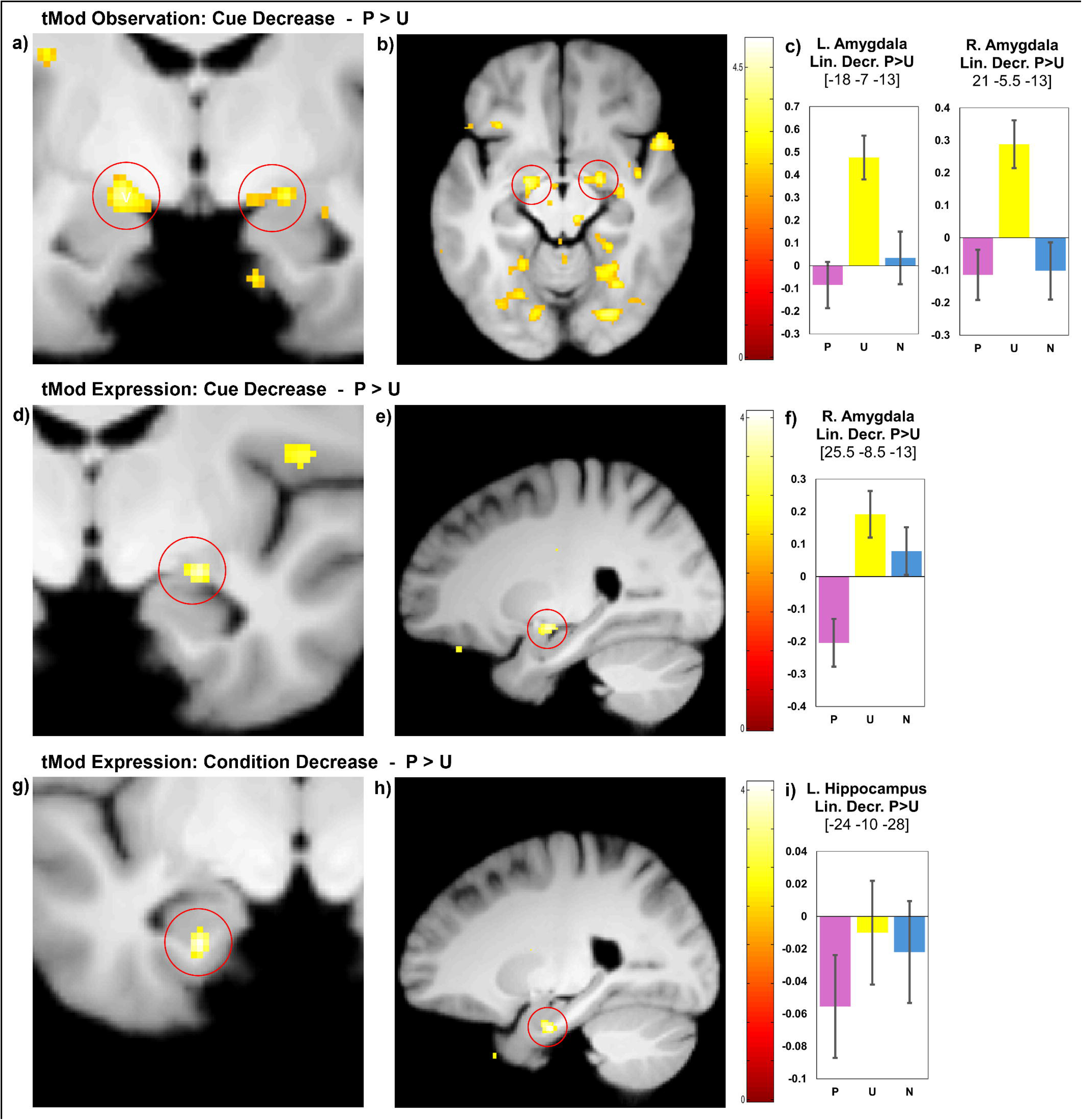
tMod cue and context. a)-c) depict the linearly decreasing contrast P > U of the cue in the observation phase. a) shows a 80×80 voxel zoom in coronary view of the bilateral centromedial amygdala clusters. b) depicts a full volume sagittal view of the same clusters. c) shows diagrams of the peak-voxel ß-estimates of both amygdala clusters. The respective clusters are highlighted with red circles in a) and b). d)-e) depict the linearly decreasing contrast P > U of the cue in the expression phase. d) shows a 80×80 voxel zoom in coronary view of the right centromedial amygdala cluster. e) depicts a full volume sagittal view of the same cluster. f) shows diagrams of the peak-voxel ß-estimates of the right amygdala cluster. The respective cluster is highlighted with a red circle in d) and e). g) – i) depict the linearly decreasing contrast P > U of the sustained context activity in the expression phase. g) shows a 80×80 voxel zoom in coronary view of the left hippocampus cluster. g) depicts a full volume sagittal view of the same cluster. i) shows diagrams of the peak-voxel ß-estimates of the left hippocampus cluster. The respective cluster is highlighted with a red circle in g) and h).

##### Condition-specific responses during threat expression

In line with finding on hippocampal-dependent contextual threat learning, we found that the hippocampus (*t* = 3.94, *df* = [1,66], MNI = [-24, -10, -28], *p(FWE)* = 0.032), in the expression phase, engaged in sustained activation across the whole trial (block regressor) in the unpredictable condition compared to the predictable condition (contrast: linear decrease U > P) (see Figure 8 g) – i)). We neither found mean (not changing) activation in the ROIs at condition-onset nor sustained activation.

### 3.4. SCR-Analysis

Analysis of SCR data was prevented by bad data quality during data acquisition. Only 13 participants had valid SCR data for both experimental phases in Experiment 2, severely limiting inferential validity. See supplement (supplementary text S2) for the analysis.

## 4. Discussion

Our results demonstrate that participants successfully learned to resolve temporal threat uncertainties through observation of others’ painful experiences, as reflected by enhanced threat expectations to predictive cues and to the onset of unpredictable conditions. Our behavioral and neural findings from observational learning are in line with insights from firsthand learning and reveal enhanced expectation of threats in the presence of predictive cues, while sustained neural responses to unpredictable conditions are elicited by the absence of temporal predictability for aversive outcomes.

On a neural level, the threat expectation when participants were confronted with predictive cues and unpredictable condition-onsets were accompanied by activation in the anterior insula and amygdala. Both neural regions were already engaged during observational threat learning, where we generally found linearly decaying activation to signals that were learned as valid predictors of threats: i.e., bilateral amygdala activity decreased to predictive cues, whereas responses increased to unpredictive cues across trials. During uncertain learning situations, sustained hippocampal activation was observed across trials. Interestingly, neural responses in the amygdala and insula, key regions involved in pain observation (Jauniaux et al., 2019; Lamm et al., 2011), were enhanced during predictable observation of painful experiences in others. This enhanced activity may reflect the additional neural resources required to resolve temporal predictability, as predictive threat cues increased activity to observed pain relative to temporally unpredictable painful experiences.

Taken together, our two experiments reveal that participants integrate temporally predictive cue-information for impending threat towards others while also identifying unpredictable threat conditions in which cues have no predictive function. Sustained responses during the unpredictable condition have been attributed to contextual fear or anxiety. These contextual sustained responses are a consequence of the absence of predictive cues (Grillon et al., 2004; Grillon & Davis, 1997), i.e. introducing a predictive cue reduces response to threatening contexts (Fonteyne et al., 2009). Our results suggest that this balance between predictive cues and unpredictable conditions also holds in observational threat learning. Similar to cue and context paradigms in firsthand learning, our results reveal that the expectation of threats is higher for the predictive cue, as compared to the predictable condition onset, and this ‘cue-prominence’ difference is larger in the predictable, as compared to the unpredictable condition. Our experimental design with moving sliders allowed for temporal anticipation of threats at the onset of the condition in the predictable (but not unpredictable) condition. Yet, the anticipation of threats was higher at the onset of the unpredictable condition (compared to the onset of the predictable condition) since in the predictable condition the informative signal for the temporal occurrence (i.e., the predictive cue) is not present. Further, our results indicated sustained expectation of threat during the unpredictable condition, when comparing the mean expectation across the onset of the condition and after passing the colored cues (see supplementary analyses).

The amygdala and hippocampus are parts of a neural network that has been found to underlie learning of firsthand threats, with studies that implied the amygdala in cued threat learning, i.e. responses of fear (Marschner et al., 2008; Naaz et al., 2019), and the hippocampus (Alvarez et al., 2011; Marschner et al., 2008) in uncertain, contextual threat. Further studies have uncovered a less clear pattern of differentiation, though (Andreatta et al., 2015; Figel et al., 2019; Herrmann et al., 2016; Hur et al., 2020). Furthermore, the role of the insular cortex in threat anticipation has been highlighted in studies that employed both temporally predictable and unpredictable threats (Alvarez et al., 2011; Geng et al., 2018; Herrmann et al., 2016; Radoman et al., 2021). Our data does not fully support a clear mechanistic separation between predictable and unpredictable threats across these brain regions. We found linearly decaying responses to predictive threat cues in the amygdala during observational learning of predictive threat cues, which aligns with reports of decaying responses to predictive cues of firsthand threats (Marschner et al., 2008). We furthermore found sustained activation in the right amygdala during observation of unpredictable threats. These findings mirror neural activation in paradigms with firsthand aversive experiences that report phasic activation of the amygdala to threat cues (Alvarez et al., 2011; Naaz et al., 2019) and unpredictable condition-onsets (Figel et al., 2019; Herrmann et al., 2016), as well as direct threat confrontation (Siminski et al., 2021). Moreover, sustained activation of the amygdala during unpredictable threat conditions has been observed in firsthand learning using electric shocks (Andreatta et al., 2015). Our analyses revealed the hippocampus to be involved in both, encoding threat cues and uncertain threat condition-onsets in observational threat learning, a seemingly contradictory finding to the proposed differential role in firsthand learning (Alvarez et al., 2011; Marschner et al., 2008), but see (Cornwell et al., 2025; Hur et al., 2020). In our study, hippocampal activation reflected the direct comparison between cue and condition-onset in the predictable vs. unpredictable conditions (i.e., the threat cue-prominence, as rated in the expression phase) already during the observation phase (as trendwise significant cluster). This role of the hippocampus in observational threat learning might be driven by an inherent uncertainty of observational learning about firsthand outcomes for the participants, i.e. even predictable observed threats being unpredictable regarding their relevance for the own outcome. Hence, the hippocampal involvement during observational learning might additionally reflect surveillance for own firsthand outcomes, rather than only temporal predictions for the demonstrator. Alternatively, hippocampal activation might also be explained merely through the temporal structuring of the trials, as hippocampal neurons have been demonstrated to be active during both spatial and temporal navigation (Bunsey & Eichenbaum, 1996; Howard & Eichenbaum, 2015; O’Keefe & Dostrovsky, 1971). Future studies that allow us to distinguish between the observer’s expectation and the observed outcomes might allow the distinction of a hippocampal function in observational learning. We found that the threat information was resolved in the left anterior hippocampus, which is in line with the functional topography of the human hippocampus, where the L1 segment is associated with fear conditioning and strongly connected to the limbic system (Robinson et al., 2015). The activation of the hippocampus to predictive cues on the other hand took place in the left posterior portion, an area primarily associated with spatial perception but with strong functional links to the left IFG involved in semantic encoding, a plausible role during learning cued threat contingencies (Robinson et al., 2015).

In line with previous research employing firsthand learning (Geng et al., 2018; Radoman et al., 2021; Shankman et al., 2014), the anterior insula was involved during observation of uncertain threat anticipation (i.e., unpredictable condition-onsets). Furthermore, the anterior insula reflected cue prominence in the expression phase, i.e., resolving condition- and cue-related threat information for unpredictable and predictable threats. This extends the finding that the anterior insular cortex computes affective responses in uncertain conditions of threat (Radoman et al., 2021).

It has been suggested for the acquisition of fear conditioning, that dynamic rather than rigid neural responses underlie the learning of threat information (Buchel, Holmes, et al., 1998; Marschner et al., 2008; Quirk et al., 1997). Consistent with previous findings (Marschner et al., 2008), we observed a linear decrease in bilateral amygdala activation towards predictive cues, while responses to unpredictive cues increased over time. This pattern persisted in the expression phase: predictive cues briefly activated the right amygdala, whereas unpredictive cues elicited sustained and increasing right amygdala activity. Similarly, the left hippocampus showed sustained activation in response to unpredictive cues. It has been argued that the amygdala (Marschner et al., 2008) as well as the anterior hippocampus (Strange et al., 1999) respond to the integration of novel information during learning. Our data fits this assumption well, with a limited amygdala response to predictable threat stimuli and a dually sustained (across length of individual trial & across trials) anterior hippocampal activation to unpredictable threats. We suggest that the predictable condition leads to rapid learning and a quick resolution of the prediction error, while unpredictability maintains continuous prediction error throughout learning (e.g. as modelled by: Rescorla et al., 1972). During the expression phase, the omission of the expected threats might further add to the decrease in threat value by extinction learning.

We found that the left BLA and anterior insula were activated during the observation of pain in others, which is in line with meta-analyses that report neural responses in these regions showing both, faces in pain, and depicting noxious stimulation of others’ extremities (Fallon et al., 2020; Jauniaux et al., 2019; Lamm et al., 2011). Interestingly, we found differences in neural responses to others’ pain, based on the learning condition: We found higher hemodynamic responses in the anterior insular cortex and the BLA to observation of others’ pain in the simpler learning condition (i.e. predictable condition), as compared to the unpredictable condition, where observed pain was not informative to learn threat signals. In the unpredictable condition, participants relied solely on social information, i.e. facial expression and gestures to infer pain, and could not predict threat towards others by non-social cues. Our results might hence imply that the observation of predictable, as compared to unpredictable pain, enhanced neural responses in the amygdala and insula as a function of learning about threat expectation and that activation of the right hippocampus might reflect the integration of the observed pain into learned predictions. Interestingly, recent research has highlighted the importance of better understanding the neural mechanisms when observed pain is integrated and when it is ignored (Wesarg-Menzel et al., 2025). A better understanding of mechanism influencing the perception of pain in others might also aid a better understanding of patient/provider interaction (Chen et al., 2019).

Firsthand cue and context learning tasks have often used startle-probes to assess whether participants learned to discriminate between cue and context related threat contingencies (Schmitz & Grillon, 2012), whereas, we used SCR and explicit expectation queries. In this context, we defined cue prominence as the difference of expectation or skin conductance measure with the cue present and at condition onset. Previous research has successfully used SCR in firsthand NPU-paradigms (Alvarez et al., 2011; Hur et al., 2020) and observational learning (Haaker, Golkar, et al., 2017; Olsson et al., 2007). Surprisingly we did not find meaningful psychophysiological reactions mirroring the explicitly given threat expectations. Potentially due to bad SCR data quality, which led to a high drop-out rate in the analysis, the assessment of psychophysiological reactions was hindered in our study.

## 5. Limitations

We established a novel paradigm and both experiments conceptually resemble each other closely but slightly differed in the rating procedure. Surprisingly, a visual comparison of the behavioral results between Experiment 1 and 2 indicated generally lower threat expectation ratings in Experiment 2. One explanation for this result could be that participants in Experiment 2 were wearing an identical cuff as seen on the demonstrator during the task and were told that it might inflict unpleasant stimulation during the expression phase. When participants did not receive any stimulation, this may have resulted in extinction learning during the expression phase. Furthermore, in both experiments, ratings collected at condition onset revealed that during unpredictable threat trials, participants were uncertain when the condition began, rather than expecting pain at condition onset (scale: -1 certainly no pain, 0 uncertain, +1 certainly pain). One might explain this fact by conceptualizing cue-prominence as an increased distance from uncertainty (i.e. 0 rating). In both U and P, expectations of pain increased towards the cue. The higher cue-prominence in P may be interpreted as resulting from a smaller reduction in uncertain expectations of pain compared to U. Furthermore, our sample size for Experiment 2 might have been small, given the signal to noise limitations of fMRI research.

## 6. Conclusion

Here, we show the behavioral effects and neural basis of observational learning in temporally predictable and unpredictable threat-situations across two experiments. Our findings demonstrate that healthy participants are able to resolve predictable and unpredictable threat contingencies by observing others’ experiences and infer expectations of self-related threat. Our results provide the neural basis for vicarious learning of cued and sustained threats, to build a basis to understand anxiety and fear, in humans as socially connected individuals. We were able to show that learning processes related to temporal certainty of threats closely mirror those from firsthand learning. On a neural level, our findings provide two novel perspectives, one that extends the neural mechanisms recruited to learn uncertain threat contingencies, and the second suggesting that such learning about threat expectations changes how we respond to observed pain of others. There is solid evidence for firsthand threat learning mechanisms being linked to anxiety (Cooper et al., 2022) and obsessive-compulsive disorders (Cooper & Dunsmoor, 2021). A better understanding of how vicarious processes contribute to threat learning might aid broadening the understanding of fear, anxiety and anxiety-related disorders.

## 7. Data availability

Raw behavioral data from both experiments and code from the fMRI analysis is available in this online repository: https://osf.io/653pr/. Wholebrain *t*-maps for all reported contrasts can be found in a neurovault online repository: https://identifiers.org/neurovault.collection:23814.

