## Supplemental Texts and Tables for "Resolving temporal threat uncertainty by observational learning involves the amygdala, hippocampus and anterior insula"

**Supplementary Text 1: Other Expectation**

A 3x2 repeated measurement ANOVA revealed a significant influence of the condition ($F = 26.685$, $df = 2$, $p < .001$), cue/context ($F = 14.936$, $df = 1$, $p < .001$) and their interaction ($F = 8.325$, $df = 2$, $p < .001$). Post-hoc t-tests of that interaction showed an increased cue-prominence in both U ($t = 4.751$, $df = 19$, $p = 0.00014$, $p_{Holm} = 0.00042$) and N ($t = 3.434$, $df = 19$, $p = 0.003$, $p_{Holm} = 0.006$) as compared to P.

**Supplementary Text 2: SCR**

13 subjects were included into the analysis after data-quality assessment in the observation phase. Only the factor cue/context significantly influenced SCR ($F = 17.309$, $df = 1$, $p = 0.001$) but not condition ($F = 1.438$, $df = 2$, $p = 0.257$) or the interaction of condition and cue/context ($F = 0.631$, $df = 2$, $p = 0.541$). A post-hoc test revealed an increased response to context onsets as compared to cues among all conditions ($t = 4.16$, $df = 12$, $p = 0.001$). Only a descriptive difference (P (mean ± SD) = 0.401 ± 0.948; U (mean ± SD) = 0.195 ± 0.794; N (mean ± SD) = 0.197 ± 0.934) among the responses to predictable pain as compared to both unpredictable ($t = 0.896$, $df = 12$, $p = 0.388$) and no pain ($t = 0.067$, $df = 12$, $p = 0.516$) was revealed.

In the expression phase 15 subjects were included after data-quality assessment. No significant effect or interaction-effect of condition or cue/context SCR was found (condition: $F = 2.285$, $df = 2$, $p = 0.120$; cue/context: $F = 3.891$, $df = 1$, $p = 0.069$; conditionXcue/context: $F = 20.083$, $df = 2$, $p = 0.921$).

**Table S1. Experiment 1:** repeated measurement ANOVA. Threat Expectation

| Cases | Sum of Squares | df | Mean Square | F | p | $\eta^2_p$ |
|---|---|---|---|---|---|---|
| Within Subjects Effects | | | | | | |
| **Condition** | 6.99 | 2 | 3.495 | 13.594 | < 0.001 | 0.475 |
| Residuals | 7.713 | 30 | 0.257 | | | |
| **Cue/Context** | 22.248 | 1 | 22.248 | 123.337 | < 0.001 | 0.892 |
| Residuals | 2.706 | 15 | 0.18 | | | |
| **Condition X Cue/Context** | 2.587 | 2 | 1.294 | 12.397 | < 0.001 | 0.452 |
| Residuals | 3.131 | 30 | 0.104 | 13.594 | | 0.475 |

**Table S1 legend.** Table S1 reports the detailed results of the behavioral effect (threat expectation) rmANOVA (3x2: Condition: P, U, N; Timepoint: cue, mean(context-onest,post-cue context)) of Experiment 1. We report sum of squares, degrees of freedom, mean square, F-values, p and an estimation of the effect sizes (partial $\eta^2$) for the main effects of condition and timepoint and the interaction effect.

**Table S2. Experiment 1:** Paired-sample t-tests. Learned cue-prominence after observation phase.

| Variable 1 | Variable 2 | df | T | p | $p_{Holm}$ | Cohen's d |
|---|---|---|---|---|---|---|
| Cue-Prominence P | Cue-Prominence N | 15 | 4.653 | 0.0003126 | 0.0009378 | 1.163 |
| Cue-Prominence U | Cue-Prominence N | 15 | 2.582 | 0.021 | 0.021 | 0.645 |
| Cue-Prominence P | Cue-Prominence U | 15 | 2.677 | 0.017 | 0.034 | 0.669 |

**Table S2 legend.** Table S2 reports the posthoc paired-sample t-tests conducted to analyze the significant interaction effect of the rmANOVA in Experiment 1. Note that the cue-prominence is here calculated as cue – mean(context-onset, post-cue context). We compare the learned cue-prominence between all three conditions. Degrees of freedom, T, p, p corrected for multiple comparisons (Bonferroni-Holm) and an estimation of effect size (Cohen's d) are reported.

**Table S3. Experiment 1:** Behavioral outcome measures

| **Variable** | **Descriptives** |
|---|---|
| Cue-Prominence P (mean ± SD) | 133.5 ± 44.1 |
| Cue-Prominence U (mean ± SD) | 101.7 ± 52.2 |
| Cue-Prominence N (mean ± SD) | 53.6 ± 55.8 |

**Table S3 legend.** Table S3 reports the descriptive results of Experiment 1. We report the calculated cue-prominence for each condition (P, U, N). Note that the cue-prominence is here calculated as cue – mean(context-onset, post-cue context).

**Table S4. Experiment 2:** repeated measurement ANOVA. Threat Expectation

| Cases | Sum of Squares | df | Mean Square | F | p | $\eta^2_p$ |
|---|---|---|---|---|---|---|
| Within Subjects Effects | | | | | | |
| **Condition** | 8.988 | 2 | 4.494 | 24.917 | <.001 | 0.531 |
| Residuals | 7.936 | 44 | 0.18 | | | |
| **Cue/Context** | 9.257 | 1 | 9.257 | 52.736 | < .001 | 0.706 |
| Residuals | 3.862 | 22 | 0.176 | | | |
| **Condition X Cue/Context** | 1.938 | 2 | 0.969 | 13.835 | < .001 | 0.386 |
| Residuals | 3.082 | 44 | 0.07 | | | |

**Table S4 legend.** Table S4 reports the detailed results of the behavioral effect (threat expectation) rmANOVA (3x2: Condition: P, U, N; Timepoint: cue, mean(context-onest,post-cue context)) of Experiment 2. We report sum of squares, degrees of freedom, mean square, F-values, p and an estimation of the effect sizes (partial $\eta^2$) for the main effects of condition and timepoint and the interaction effect.

**Table S5. Experiment 2:** Paired-sample t-tests. Learned cue-prominence after observation phase.

| **Variable 1** | **Variable 2** | **df** | **T** | **p** | **$p_{Holm}$** | **Cohen's d** |
|---|---|---|---|---|---|---|
| Cue-Prominence P | Cue-Prominence N | 22 | 4.497 | 0.0001793 | 0.0005379 | 0.352 |
| Cue-Prominence U | Cue-Prominence N | 22 | 3.671 | 0.001 | 0.002 | 0.288 |
| Cue-Prominence P | Cue-Prominence U | 22 | 1.631 | 0.117 | 0.117 | 0.185 |

**Table S5 legend.** Table S5 reports the posthoc paired-sample t-tests conducted to analyze the significant interaction effect of the rmANOVA in Experiment 2. Note that the cue-prominence is here calculated as cue – mean(context-onset, post-cue context). We compare the learned cue-prominence between all three conditions. Degrees of freedom, T, p, p corrected for multiple comparisons (Bonferroni-Holm) and an estimation of effect size (Cohen's d) are reported.

**Table S6. Experiment 2:** Behavioral outcome measures

| Variable | Descriptives |
|---|---|
| Cue-Prominence P (mean ± SD) | 75.5 ± 50.5 |
| Cue-Prominence U (mean ± SD) | 70.5 ± 52.1 |
| Cue-Prominence N (mean ± SD) | 19.4 ± 32.5 |

**Table S6 legend.** Table S6 reports the descriptive results of Experiment 2. We report the calculated cue-prominence for each condition (P, U, N). Note that the cue-prominence is here calculated as cue – mean(context-onset, post-cue context).
